# *Dugesia japonica* is the best suited of three planarian species for high-throughput toxicology screening

**DOI:** 10.1101/2020.01.23.917047

**Authors:** Danielle Ireland, Veronica Bochenek, Daniel Chaiken, Christina Rabeler, Sumi Onoe, Ameet Soni, Eva-Maria S. Collins

## Abstract

High-throughput screening (HTS) using new approach methods is revolutionizing toxicology. Asexual freshwater planarians are a promising invertebrate model for neurotoxicity HTS because their diverse behaviors can be used as quantitative readouts of neuronal function. Currently, three planarian species are commonly used in toxicology research: *Dugesia japonica*, *Schmidtea mediterranea*, and *Girardia tigrina*. However, only *D. japonica* has been demonstrated to be suitable for HTS. Here, we assess the two other species for HTS suitability by direct comparison with *D. japonica*. Through quantitative assessments of morphology and multiple behaviors, we assayed the effects of 4 common solvents (DMSO, ethanol, methanol, ethyl acetate) and a negative control (sorbitol) on neurodevelopment. Each chemical was screened blind at 5 concentrations at two time points over a twelve-day period. We obtained two main results: First, *G. tigrina* and *S. mediterranea* planarians showed significantly reduced movement compared to *D. japonica* under HTS conditions, due to decreased health over time and lack of movement under red lighting, respectively. This made it difficult to obtain meaningful readouts from these species. Second, we observed species differences in sensitivity to the solvents, suggesting that care must be taken when extrapolating chemical effects across planarian species. Overall, our data show that *D. japonica* is best suited for behavioral HTS given the limitations of the other species. Standardizing which planarian species is used in neurotoxicity screening will facilitate data comparisons across research groups and accelerate the application of this promising invertebrate system for first-tier chemical HTS, helping streamline toxicology testing.

## Introduction

Toxicology is currently undergoing a paradigm shift, focusing considerable effort on replacing, reducing, and refining (3Rs) vertebrate animal testing. This change has been driven by the high cost, low throughput, and questionable relevance of traditional mammalian guideline tests used for regulatory decisions. This is especially true for assessing developmental neurotoxicity (DNT) (Tsuji and Crofton, 2012). New approach methods which are amenable to economical high-throughput screening (HTS), including *in silico* modeling, *in vitro* models, and invertebrate systems, promise to fill the gap, alone or as part of a test battery (Fritsche et al., 2018; Lein et al., 2005; Thomas et al., 2019). A recent directive from the Environmental Protection Agency (EPA) details a plan to stop all funding of mammalian testing by 2035 (Wheeler, 2019). This directive reinforces the agency’s previous commitment to reduce vertebrate testing for chemicals regulated under the Toxic Substances Control Act through integration of new approach methods into regulatory decisions (US EPA, 2018). To achieve this challenging goal, an increased effort is necessary to validate these new approach methods to ensure sensitivity, robustness, and relevance, and standardize best testing practices (Bal-Price et al., 2018; Crofton et al., 2011). Common test standards for a particular model system are essential for meaningful direct comparisons of data across laboratories and ultimately will build the basis for the development of the necessary regulatory guidelines.

We have developed the asexual freshwater planarian *Dugesia japonica* as a promising new invertebrate model for high-throughput neurotoxicity and DNT screening (Hagstrom et al., 2016, 2015; Zhang et al., 2019a, 2019b). We have shown that it possesses comparable sensitivity to more established new approach methods and is predictive of mammalian DNT (Hagstrom et al., 2019, 2015; Zhang et al., 2019a, 2019b). The key advantage of the planarian system is its sufficiently complex behavioral repertoire which enables distinct behaviors to be used as a multifaceted quantitative readout of neuronal function (Hagstrom et al., 2019; Zhang et al., 2019a, 2019b). The planarian nervous system is of medium size (∼10,000 neurons), possessing >95% gene homology and sharing most of the same neurotransmitters and neuronal cell types as the mammalian brain (Buttarelli et al., 2008; Mineta et al., 2003; Ross et al., 2017). Thus, the planarian system allows for mechanistic insights into how different cells and pathways control specific behaviors (Birkholz and Beane, 2017; Currie and Pearson, 2013; Inoue et al., 2015, 2014; Nishimura et al., 2010, 2008; Pearce et al., 2017; Sabry et al., 2019; Zhang et al., 2019b). Because planarians are simultaneously amenable to high-throughput screening (HTS), they are a promising alternative neurotoxicology model. We and others have recently reviewed the benefits and limitations of planarians for toxicology, particularly neurotoxicity and DNT (Hagstrom et al., 2016; Wu and Li, 2018; Zhang et al., 2019a).

Our previous work demonstrated the potential of *D. japonica* as an invertebrate model for neurotoxicity and DNT studies and demonstrated the reliability and robustness of our screening methodology (Hagstrom et al., 2015; Zhang et al., 2019a, 2019b). However, since other research groups have used other planarian species and other, generally low-throughput and small scale, screening methods, it is difficult to compare results or standardize testing conditions (Hagstrom et al., 2016; Wu and Li, 2018; Zhang et al., 2019a). The two most common planarian species that have been used in toxicology studies besides *D. japonica* are *Girardia tigrina,* formerly *Dugesia tigrina,* (Byrne, 2018; Córdova López et al., 2019; Knakievicz and Ferreira, 2008; Moustakas et al., 2015; Ramakrishnan and DeSaer, 2011) and *Schmidtea mediterranea* (Lowe et al., 2015; Plusquin et al., 2012; Poirier et al., 2017; Stevens et al., 2014; Tran et al., 2019). Of these three, *S. mediterranea* is the most popular planarian species for molecular studies because its annotated genome is readily available (Grohme et al., 2018; Robb et al., 2008; Rozanski et al., 2019), whereas only a draft genome exists for *D. japonica* (An et al., 2018). Transcriptomes are available for all three species (Rozanski et al., 2019; Wheeler et al., 2015). Genomic studies have been hindered in *D. japonica* and *G. tigrina* because of the larger size (2n=16, compared to 2n=8 in *S. mediterranea*), mixoploidy, and abundance of repetitive, transposable elements in the genomes of these species (An et al., 2018; Benazzi, 1993; Garcia-Fernandez et al., 1995; Hoshino et al., 1991; Wheeler et al., 2015). Comparatively, *G. tigrina* is the least well characterized, but is commercially available and has thus found widespread use across research laboratories and schools. *G. tigrina* has been largely utilized for its characteristic head morphology (auricles), which facilitates scoring of morphological head abnormalities and regeneration defects (Córdova López et al., 2019; Knakievicz and Ferreira, 2008).

We have previously found that there are significant differences in terms of growth and reproductive strategies in the laboratory among these three species (Carter et al., 2015). Most relevant in respect to HTS suitability are our findings that *G. tigrina* and *S. mediterranea* are more sensitive to water conditions than *D. japonica* (Carter et al., 2015), which could be problematic when these species are stored in small volumes for extended periods of time, such as during HTS in multi-well plates.

In the context of toxicology screens, only *D. japonica* has so far been tested and demonstrated to be a suitable HTS system (Zhang et al., 2019a, 2019b), because the same repertoire of behaviors which can be observed in low-throughput experiments are reproducible in a HTS setting (a sealed 48-well plate, with 1 planarian per 200 µl of solution per well) (Hagstrom et al., 2015; Zhang et al., 2019a) and specimen can be recovered from the HTS setup without obvious long-term negative health effects.

Thus, we aim to evaluate two criteria: 1) which species is the best suited for HTS conditions and 2) how sensitive the different species are to solvents commonly used in toxicology. To directly compare the suitability of these three planarian species for HTS, we utilized our custom robotic screening platform because it was demonstrated to be reliable and robust (Zhang et al., 2019b). On this automated platform, chemicals are screened in a 48-well plate, testing 5 concentrations along with a solvent control for n=8 planarians (1/well) per condition and experiment. Planarian morphology and behaviors are assayed and quantified at Days 7 and 12 of neurodevelopment/exposure (Zhang et al., 2019a). Regeneration occurs on similar time scales for the three species, allowing comparisons to be made using the same time points.

The use of solvents is often necessary for chemical testing, particularly for aqueous solutions; thus, it is important to assess the potential toxicity of relevant solvent concentrations to ensure this does not interfere with assessment of test chemicals. Therefore, we assayed 4 common solvents (dimethyl sulfoxide (DMSO), ethanol, methanol, ethyl acetate) and a negative control (sorbitol) at concentrations previously determined to be sublethal in *D. japonica* (Hagstrom et al., 2015; Zhang et al., 2019a).

Unexpectedly, we found that under these HTS test conditions, *S. mediterranea* and *G. tigrina* exhibited limited motility, hindering our ability to evaluate meaningful morphological and behavioral defects in these species. In addition, these species tended to be more sensitive to solvent toxicity than *D. japonica*. For example, significant lethality was observed in methanol in *S. mediterranea* and *G. tigrina,* but only behavioral defects were found in *D. japonica* at the same concentrations. Together, our data show that *D. japonica* performs the best under the experimental constraints required for HTS and thus is the species of choice for planarian HTS.

## Material and Methods

### Specimen

Asexual *D. japonica*, *G. tigrina*, and *S. mediterranea* freshwater planarians were cultivated using standard protocols. *D. japonica* and *S. mediterranea* planarians were from established lab cultures. *G. tigrina* planarians were purchased from Ward’s Science (Rochester, NY, USA) and thus is it unknown how long this population has been reared under laboratory conditions. *S. mediterranea* were kept in 1X Montjüic salts (Cebrià and Newmark, 2005). *D. japonica* and *G. tigrina* were stored in dilute (0.5 g/L) Instant Ocean Salts (IO) (Spectrum Brands, Blacksburg, VA, USA). For simplicity, “planarian water” will refer to the respective water used for each species. Planarians were stored in tupperware containers at 20°C in a temperature-controlled Panasonic incubator in the dark when not used for experiments. The animals were fed organic chicken or beef liver 1-2 times per week and cleaned twice per week (Dunkel et al., 2011). Liver was purchased frozen from a local farm, thawed, cut into small pieces and aliquoted. Aliquots were stored at −20 °C for up to 6 months before use. For experiments, we randomly selected similarly sized, intact planarians that were starved 5-7 days prior to experiment onset. On Day 1, selected specimens were amputated pre-pharyngeally via an ethanol-sterilized razor blade.

### Chemical Preparation

Table 1 summarizes the details on the chemicals and concentrations that were used. The highest tested concentration of each solvent was chosen to be sublethal to *D. japonica*, as determined from previous experiments (Hagstrom et al., 2015) and by preliminary testing. Stocks of all chemicals were prepared in IO water at 10x of the highest tested concentration. Experimental concentrations were made using 2-fold serial dilutions in IO water. D-sorbitol (D-glucitol) served as a negative control (Zhang et al., 2018) and was prepared using serial half-log dilutions in IO water. All dilutions were made and used fresh on Day 1 of the assay.

**Table 1.** Tested solvents and their experimental concentrations.

| Chemical Name | CAS# | Supplier | Purity (%) | Tested Concentrations |
| --- | --- | --- | --- | --- |
| Ethanol | 64-17-5 | Greenfield Global | (ACS): 99.98%,<br>(USP): 99.99%. | 1, 0.5, 0.1, 0.05, 0.01 (% v/v) |
| Methanol | 67-56-1 | Sigma Aldrich | 99.9 | 3.2, 1.6, 0.8, 0.4, 0.2 (% v/v) |
| DMSO | 67-68-5 | Sigma Aldrich | 99.9 | 1, 0.5, 0.1, 0.05, 0.01 (% v/v) |
| Ethyl Acetate | 141-78-6 | Sigma Aldrich | 99.8 | 0.04, 0.02, 0.01, 0.005, 0.0025 (% v/v) |
| D-sorbitol | 50-70-4 | Sigma Aldrich | 99 | 100, 31.6, 10, 3, 1 (μM) |

### Exposure set-up

For every chemical concentration and planarian species, 3 technical replicates of n=8 (n=24 in total) developing/regenerating planarians were assayed in independent screening plates, using independent chemical preparations. Screening plates were prepared as described in (Zhang et al., 2019a). In brief, on Day 1 of the screen, planarians were decapitated, and their tails were randomly placed in separate wells of a 48-well plate (Genesee Scientific, San Diego, CA) (1 worm/well) containing 180 µl of planarian water. 20 µl of 10x stocks of the respective chemicals or the vehicle control were added to the screening plates within 3 hours following amputation. For each chemical and replicate, 1 screening plate was prepared such that the 5 test concentrations and 1 vehicle control (planarian water) were contained within the plate (one condition per row). The concentration pattern in each plate was shifted down 2 rows in each replicate to control for edge effects (Zhang et al., 2019a). Plates were sealed with ThermalSeal RTS sealing film (Excel Scientific, Victorville, CA) and stored in the dark at room temperature for the duration of screening (12 days). The plates were moved to the screening platform only when screened on Day 7 and Day 12. Chemical solutions were not replaced over the course of the screening period.

### Planarian motility experiments

To test why *G. tigrina* and *S. mediterranea* planarians showed limited motility under the HTS screening conditions, we set up 48-well plates as described above using regenerating or intact planarians of each species. For regenerating planarian tests, we first screened the initial intact worms in a 48-well plate within 30 min of plate setup. The planarians were then amputated as described above, allowed to regenerate in petri dishes, and screened in 48-well plates again on Days 7 and 12. For intact planarian tests, the intact planarians were first confirmed to show normal locomotion under white light conditions in a petri dish. The planarians were then loaded into 48-well plates, which were sealed as described above. The 48-well plates were screened within 30 min of plate setup and again on Day 7 and Day 12. In addition, we tested the behavior of *S. mediterranea* planarians under different lighting conditions. Specifically, we compared their locomotion and thermotaxis behavior under red light conditions, as used in our HTS setup, with those under white light conditions, by laterally adding white light illumination to the assay station. For *G. tigrina* low throughput thermotaxis experiments, we used a custom peltier to assay 6 wells of a 6-well plate simultaneously (3 planarians/well). Ambient red lighting from an electroluminescence strip was used. Wells were filled with 3 ml IO water/well. Three wells contained *D. japonica* planarians (as experimental controls for the gradient) and 3 wells *G. tigrina*. Plate loading was rotated between triplicate experiments to account for any variability in gradient strength across the peltier. Plates were recorded for 2 min without and then 4 min with the gradient on.

### Screening platform

Our custom screening platform consists of a commercial robotic microplate handler (Hudson Robotics, Springfield Township, NJ), two custom-built imaging systems, and multiple assay stations as described in detail in (Zhang et al., 2019a). The imaging systems, assay stations, and plate handler were controlled by a computer. Image analysis was performed using custom MATLAB or Python scripts. In addition to the assays performed in (Zhang et al., 2019a), we have expanded the platform in the following ways (described in detail below): 1) modification of the phototaxis assay to increase the resting period before the blue light stimulus, 2) modification of the scrunching assay to capture differences in the timing of reaction, and 3) addition of an automated “stickiness” assay. Moreover, analysis of the morphology/regeneration assay was expanded to also detect body shape changes.

First, the timing of the phototaxis assay was modified to increase the resting time in the red light (dark cycle) to 2 minutes before a 1 min blue light stimulation (light cycle), though only the activity in the last minute in the dark cycle was analyzed. The increased time in the dark cycle allowed the planarians to acclimate and settle before the blue light stimulus. The phototactic response was quantified by calculating the difference of the average speed in the blue light cycle to that in the preceding 1 min of the dark cycle (Zhang et al., 2019a). Dead planarians were discarded from the analysis.

Second, the scrunching assay was modified to allow for a dynamic analysis of noxious heat sensing as we previously found that some chemicals interfere with the rate of reaction to noxious heat (Hagstrom et al., 2018). Thus, the rate of heating of the peltier was modified to allow for a more gradual ramping up in temperature. In addition to the binary scoring of scrunching, two new endpoints were added to this assay to evaluate 1) the rate of reaction and 2) the strength of reaction to the noxious heat. Similar to (Hagstrom et al., 2018), the center of mass (COM) of each planarian was tracked over the course of the experiment and the displacement (scaled by body length) of each worm across 6 second intervals was calculated in MATLAB. The mean displacement for every 30 second bin was then calculated. We previously found that under similar, low-throughput conditions, wild-type *D. japonica* exposed to noxious heat exhibit frequent turns and decreased movement followed by eventual paralysis (Hagstrom et al., 2018). Thus, the assay was separated into phases: 1) the initial dynamic reaction and 2) the persistent decreased movement once the reaction stabilized (Supplementary Figure 1). During the initial reaction phase, the mean displacement of control *D. japonica* planarians generally decreases over time. Therefore, the rate of reaction was quantified as the slope of mean displacement for the first 2.5 minutes of the assay, which is typically negative for control *D. japonica*. Of note, quantification of this endpoint required that the planarian moved within at least three (of the five total) 30 second intervals in the first 2.5 minutes of the assay. During the second half of the assay, *D. japonica* planarians tend to become mostly immobile but may still move their heads or wiggle in place, resulting in small displacements. Therefore, the strength of the reaction was quantified as the mean of mean displacement during minutes 3-5 of the assay (Supplementary Figure 1).

Third, we introduced a new assay, which we named the “stickiness assay” since it quantifies the worm’s tendency to stick/adhere to the substrate. This new assay is a high-throughput implementation of a previous low-throughput endpoint which we have shown is correlated with mucus production (Hagstrom et al., 2018; Malinowski et al., 2017). A microplate orbital shaker (Big Bear Automation, Santa Clara, CA) was used to shake the screening plates and thus create controlled water flow within each well to unstick the planarians from the bottom of the plate well. Different rotation speeds for regenerating planarians at Day 7 and 12 were chosen based on preliminary testing to achieve a reproducible majority fraction of wild-type *D. japonica* planarians to unstick. This intermediate unsticking capacity was chosen to be able to detect both an increase or decrease in planarian “stickiness”. Day 7 was observed as the relatively stickiest time-point, potentially due to locally increased secretion of mucus because the worms are less motile during regeneration. At Day 7, the plates were shaken for 3 seconds at 1017 revolutions per minute (rpm), whereas at Day 12, the plates were shaken for 3 seconds at 665 rpm. The plate was imaged from above by a USB3 camera (FLIR Systems Inc., Wilsonville, OR) mounted on a ring stand and imaged at 8 frames per second (fps).

Each worm was scored as either “unstuck” (defined as being displaced by the water flow and floating in the well) or “stuck” (defined as worms which did not float during the whole plate shaking session) using a custom script written in Python using functions from the Scikit-Image (Van Der Walt et al., 2014) library. The script analyzes a series of 50 frames of a plate, with approximately the first 30 frames showing the plate shaking and applies four major steps. First, the plate is cropped, registered, and segmented into 48 wells for each frame. Li thresholding (Li and Tam, 1998) is used to segment the plate from the background. For each well, candidate worms are identified across each shaking frame by removing the lightest 80 percent of pixels in each well (since the worms are the darkest object) and using three successive rounds of Otsu segmentation (Otsu, 1979). A morphological opening is applied to join close objects and a morphological closing is applied to remove small objects. The area and COM of each detected object is calculated. For the first frame of the shaking well, the largest object is identified as the worm; for each following frame, the nearest segmented object is the worm. Third, the total movement of the worm is measured by summing the distance between the weighted COM of the identified worm in each frame. Weights are determined by the ratio of the areas of the identified worms in adjacent frames, accounting for uncertainty by down-weighting movements where the frames have disagreements about the size of the worm. Fourth, stickiness is classified based on the tracked movements. For wells where a worm is detected, the well is marked as “stuck” when there is a mean of fewer than five units of movement of COM per frame or marked as “unstuck” when there are greater than five units of movement per frame. Wells for which a worm is never detected are marked as uncertain. In wells with multiple worms detected, as a result of the planarians undergoing fission during the screen duration, the stickiness of the larger worm is detected. Parameters for this script were determined using an independent test set of images of *D. japonica* planarians.

As a quality control check of this new methodology, we quantified the accuracy of the automated analysis by comparing to manual scoring (Supplementary Figure 2). Worms which were dead, were not visible by eye, or which were flagged as “unsure” by the automated analysis were excluded from the accuracy calculations. For *D. japonica* planarians, the automated analysis had an average accuracy of 84 and 88 % for Day 7 and 12, respectively. The automated analysis had slightly reduced accuracy for *S. mediterranea* and *G. tigrina,* ranging from 68-77% for the two days, due to an underestimation of stickiness. Thus, while there is room for improvement, the automated analysis works reasonably well for classifying stickiness in *D. japonica*.

Additionally, in the morphology assay, different body shapes were classified for each alive planarian, including normal body shape, general sickness (lesions, loss of pigment, head regression), contraction, curled up or C-shape, corkscrew-like, and pharynx extrusion. Of note, one planarian could be classified as having multiple body shapes, for example, C-shape and pharynx extrusion.

All assays were performed in the following order, whereby the notation in brackets indicates on which day(s) the assay was performed: phototaxis (D7/D12), unstimulated locomotion (D7/D12), stickiness (D7/D12), lethality/fission/morphology (D7/D12), eye regeneration (D7), thermotaxis (D7/D12), and scrunching (D12). Any data analysis which had to be cross-checked manually was performed blinded by a single investigator, who was not given the chemical identity of the plates.

### Statistical Analysis

Statistical testing was performed on compiled data from the triplicate runs. For all endpoints, comparisons were made between the test population and the internal set of controls for that chemical. For lethality, eye regeneration, body shape morphology, stickiness, and scrunching endpoints, a one-tailed Fisher’s exact test was used. For thermotaxis, phototaxis, noxious heat sensing, and unstimulated behavioral endpoints, Tukey’s interquartile test was first used to remove any outliers, with at most 5% of the data removed. A non-parametric one-tailed Mann Whitney U-test was used to determine significant effects in thermotaxis. For unstimulated behavior endpoints (speed and fraction of time resting) and noxious heat endpoints (rate and strength of reaction), Lilliefors test was first used to test the normality of the samples. Thus, we performed either a parametric two-tailed t-test or a nonparametric two-tailed Mann-Whitney U-test depending on whether the sample distributions were normal or not, respectively.

Statistical significance was determined as instances where the p-value was less than 0.05. When a single plate in the triplicates was responsible for designating a “hit,” the triplicate was considered inconsistent and excluded as a hit. The lowest observed effect level was determined as the lowest concentration designated as a statistically significant hit. All data are available upon request.

## Results

### Need for standardization of planarian species used in toxicological studies

The use of freshwater planarians in toxicological studies has been increasing in recent years from only a handful of papers published annually prior to 2000 to approximately 20 papers published annually in recent years (Wu and Li, 2018). Three planarian species (*D. japonica*, *S. mediterranea* and *G. tigrina*) have emerged as the most popular planarian models used for toxicological studies (Figure 1), because they are widely available and have published genomes (Grohme et al., 2018; Robb et al., 2008; Rozanski et al., 2019) or transcriptomes (Rozanski et al., 2019; Wheeler et al., 2015). In addition, stereotypical behaviors in these species have been characterized and employed as readouts for neuronal function, albeit to differing extents and with differing levels of throughput (Supplementary Table 1).

**Figure 1.**
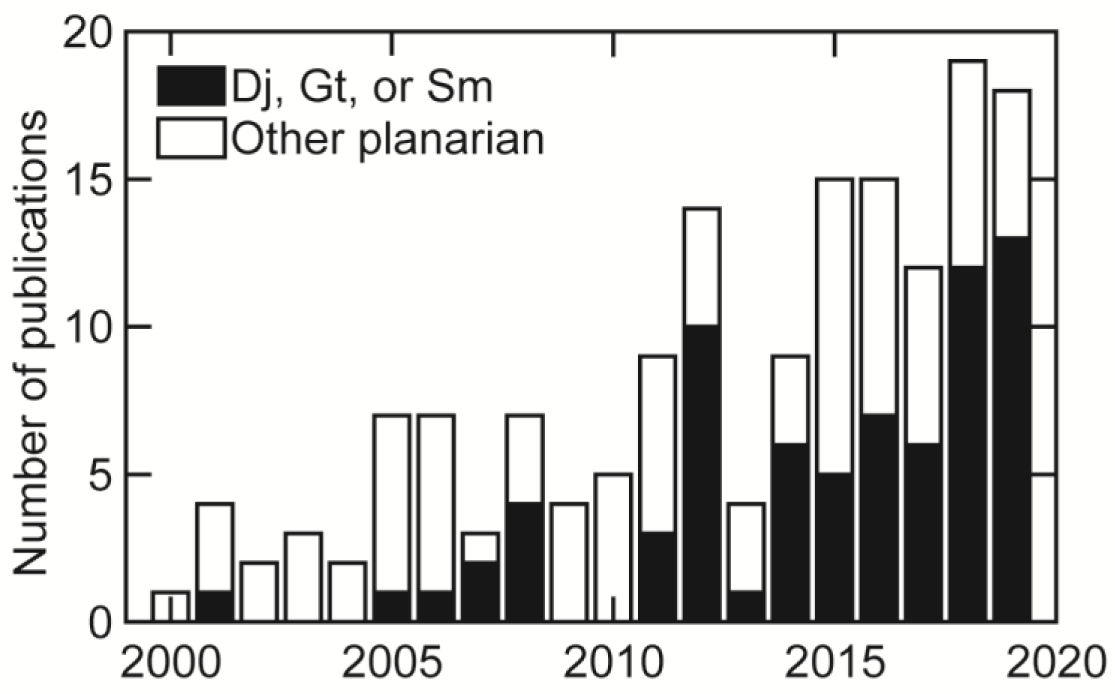
Number of journal articles reporting toxicological effects on planarian species over time. Literature search was conducted using PubMed with the following keywords: (((planarian OR flatworm) NOT marine NOT parasitic NOT Schistosoma) AND (toxic) NOT review) and (((Dugesia japonica) OR (Schmidtea mediterranea) OR ((Girardia OR Dugesia) tigrina)) AND (toxic) NOT review).

To directly compare the performance of these 3 popular species, we screened for potential morphological and behavioral effects of 4 common solvents (DMSO, ethanol, methanol, ethyl acetate) and 1 negative control (sorbitol) on regenerating *D. japonica*, *S. mediterranea,* and *G. tigrina* planarians using a robotic screening platform (Zhang et al., 2019a). We evaluated 1) the suitability of each planarian species for HTS by analyzing their performance in automated assays and 2) the sensitivity of each species to solvents commonly used in toxicology. For simplicity, we first report on the overall performance of each species, using data from control populations in the different morphological and behavioral assays, as this performance directly impacts our ability to assess chemical sensitivity in the different species.

### Lethality, body shape, and eye regeneration

As in previous screens (Zhang et al., 2019a, 2019b), control *D. japonica* exhibited very little background lethality. In contrast, significant lethality was observed in both *S. mediterranea* and *G. tigrina* control populations at Day 12, with approximately 14% and 7% lethality, respectively, (p-values: 0.015 [*S. mediterranea*] and 0.0027 [*G. tigrina*] compared to *D. japonica* using Fisher’s exact test) (Figure 2A). Death can also occur by “suicide” wherein planarians leave the water and subsequently dry out (Zhang et al., 2019a). We excluded suicides from our lethality statistics because the mechanism causing death is different. A significant number of suicides (9%) were observed in *S. mediterranea* control planarians but were not observed in the control populations of the other two species.

**Figure 2.**
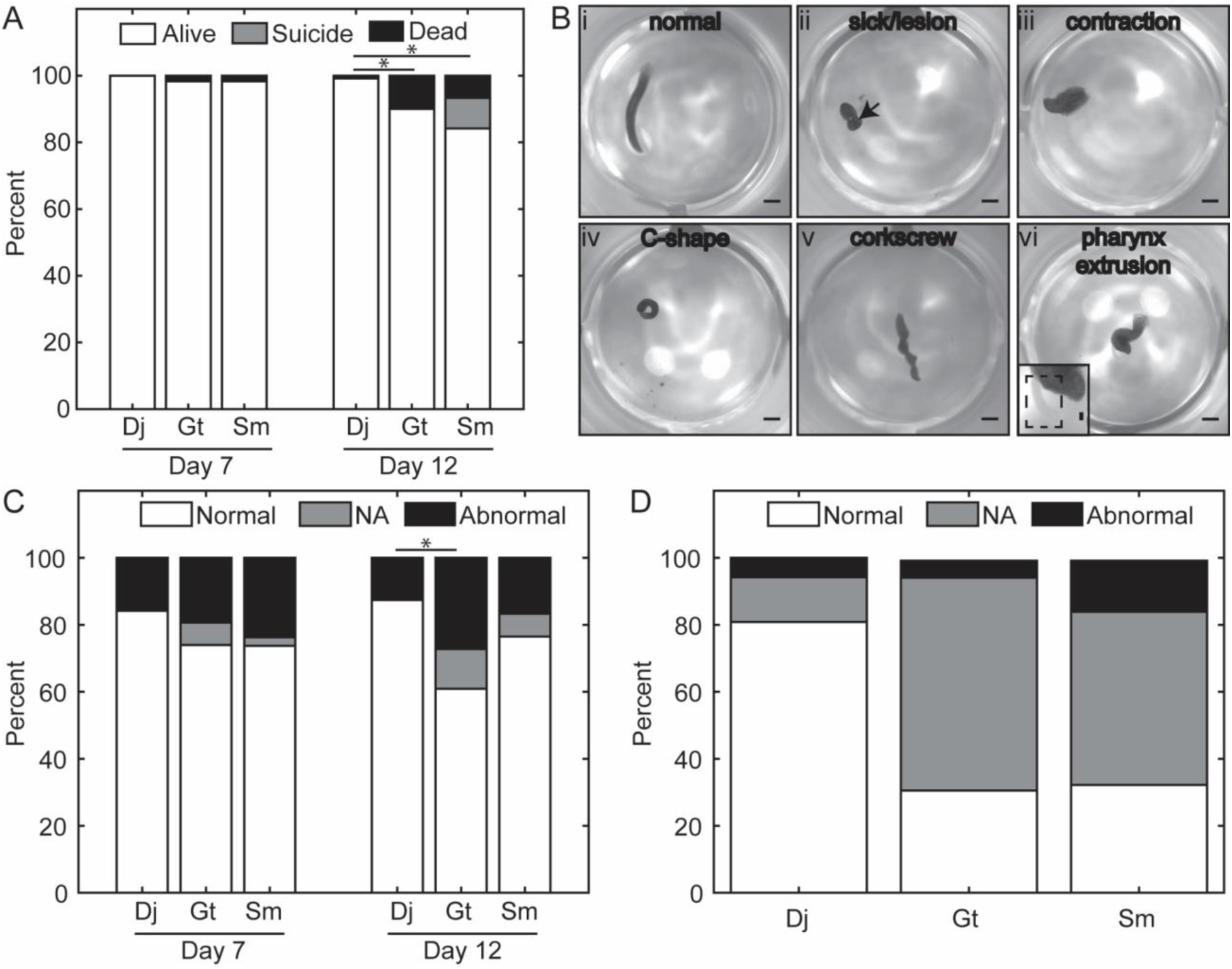
Lethality and morphology are compromised in *S. mediterranea* and *G. tigrina* controls. A) Percentage of control planarians which were alive, dead, or committed “suicide” for each species on Days 7 and 12. n=120. * indicates statistical significance with p-values < 0.05 as determined by a Fisher’s exact test. B) Examples of normal and abnormal body shapes. i) Normal planarian, ii) sick, iii) contracted, iv) curled or C-shape, v) corkscrew, vi) pharynx extrusion. Arrow points to a lesion. Inset shows pharynx protruding outside the planarian body. Scale: 1 mm. C) Percentage of alive control planarians demonstrating abnormal or normal body shapes in each species at Days 7 and 12. NA indicates the planarians could not be analyzed. * indicates statistical significance with p-values < 0.05 as determined by a Fisher’s exact test. D) Percentage of alive control planarians in each species which showed normal (2 eyes) or abnormal (0 or 1 eye) eye regeneration at Day 7. NA indicates the planarians could not be analyzed.

Planarians can exhibit a variety of abnormal morphologies and body shapes, including signs of general sickness (e.g. lesions, loss of pigment, or head regression), contraction, being curled up or C-shape, corkscrew-like, and displaying pharynx extrusion (Figure 2B). Some body shapes have been associated with disturbances to specific neurotransmitter systems (Buttarelli et al., 2008; Passarelli et al., 1999), making body shape a potentially sensitive readout for neurotoxicity. In all three species, some abnormal body shapes were observed in control populations (Figure 2C). Generally, more abnormalities were observed at Day 7 than Day 12. At both Day 7 and Day 12, the most prominent abnormal body shape in all species was contraction. At Day 7, approximately 16% of *D. japonica* controls exhibited some abnormal body shape, whereas in *S. mediterranea* and *G. tigrina* abnormal body shapes were found in 24% and 19% of controls, respectively, though these differences were not statistically significant. Only control *G. tigrina* showed greater abnormal body shapes at Day 12 than at Day 7, which was significantly greater than *D. japonica* at Day 12 (p-value< 0.001).

Analysis of lethality and body shape in *S. mediterranea* and *G. tigrina* was hindered by the fact that many of these planarians were sitting on or at the well edge for the entire morphology assay. For many of these, assessments of lethality and body shape had to be manually cross-checked by observing the animals in the other assays, which is not a viable approach for HTS. Particularly for body shape, even manual cross-checks were insufficient to determine the morphology of some planarians. In these cases, the planarians were scored as “NA” for not analyzable and were excluded from further analysis (Figure 2C). Eye regeneration was also assessed through high resolution imaging of individual wells to discern whether the planarian has regained both eyes (normal condition) or not (abnormal) (Zhang et al., 2019a). However, due to the limited visibility of many of the *S. mediterranea* and *G. tigrina* planarians in this assay, it was impossible to assess eye regeneration status in many of the control worms (Figure 2D). This led to very low sample sizes (Supplementary Table 2), decreasing the statistical power of this assay for these species.

### Stickiness

Normal planarian locomotion relies on cilia beating in a layer of secreted mucus (Martin, 1978; Rompolas et al., 2010). We previously found that increased mucus production is correlated with increased “stickiness” of the worm, which can be assessed by evaluating how easily the planarian is dislodged from its substrate, and that certain chemicals can increase planarian stickiness (Hagstrom et al., 2018; Malinowski et al., 2017). We have automated this originally low-throughput assay (Hagstrom et al., 2018; Malinowski et al., 2017), which now relies on shaking of the screening plate to create controlled water flow with the potential to unstick the planarian from the bottom of the well (Figure 3A-B).

**Figure 3.**
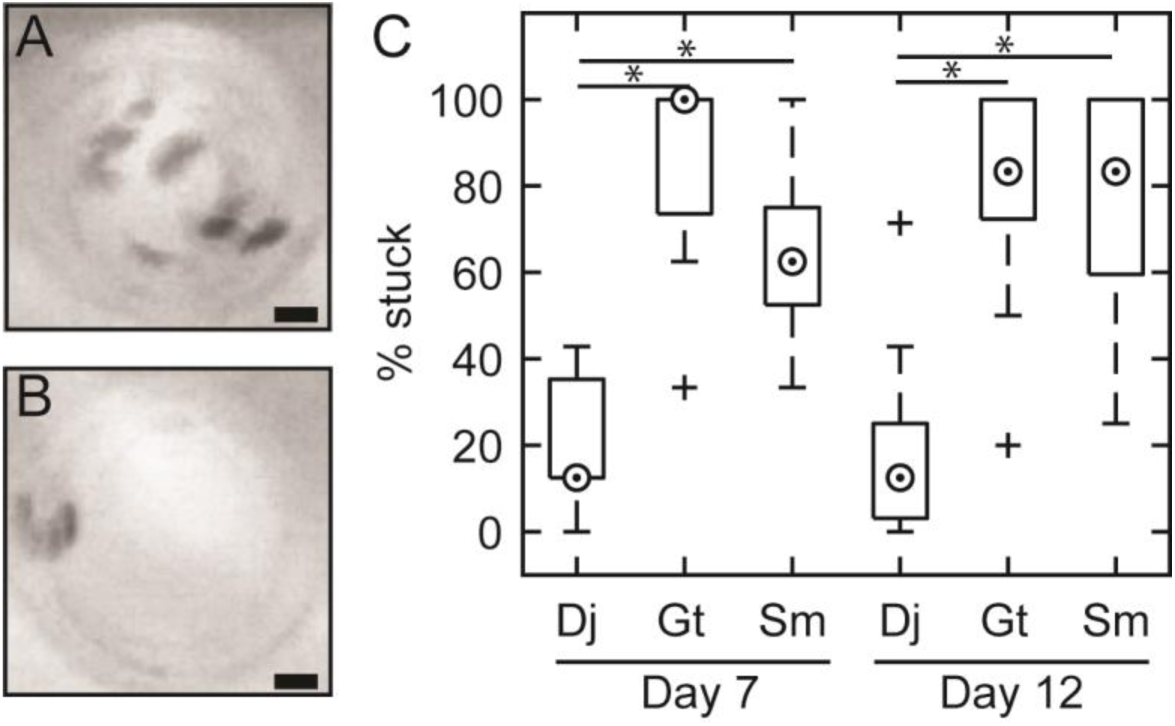
Overview of stickiness assay. (A-B) Minimum intensity projections of the shaking phase of the stickiness assay showing an (A) unstuck or (B) stuck planarian. Scale bars: 2 mm. C) Boxplot of the percent control planarians stuck in each replicate plate (n=8 per data point, n=15 data points per condition) as determined using manual analysis. Medians are shown as a dot in a circle, outliers are shown as crosses. * indicates statistical significance with p-values < 0.05 using the Mann-Whitney U-Test.

The shaking parameters were defined such that a reproducible majority of control *D. japonica* planarians would be unstuck, allowing for identification of conditions that caused either increased or decreased stickiness. We found that the other two species were significantly stickier than *D. japonica*, as the majority of controls in both *S. mediterranea* and *G. tigrina* were still stuck after shaking, and exhibited larger plate-to-plate variability (p-values: 1.1 × 10^-5^, 2.6 × 10^-5^ [*S. mediterranea* Day 7 and 12] and 4.9 × 10^-6^, 1.5 × 10^-5^ [*G. tigrina* Day 7 and 12] compared to *D. japonica* using Fisher’s exact test) (Figure 3C).

### S. mediterranea and G. tigrina show decreased motility under HTS conditions

Next, we assayed the planarians’ unstimulated locomotion by quantifying speed and the fraction of time resting. Unexpectedly, we found that both *S. mediterranea* and *G. tigrina* controls had significantly decreased motility evidenced by the large fraction of time spent resting (p-values: 5.8 × 10^-31^, 2.1 × 10^-23^ [*S. mediterranea* Day 7 and 12] and 1.9 x10^-15^, 3.0 × 10^-4^ [*G. tigrina* Day 7 and 12] compared to *D. japonica* using a two-tailed student’s t-test), (Figure 4A). Because gliding speed is only calculated for animals that glide for at least 10 continuous frames (out of 900 total), this large amount of resting greatly reduced the sample size for this endpoint (Supplementary Table 2). This effect was more pronounced in *S. mediterranea* and in Day 7 for both *S. mediterranea* and *G. tigrina*. Moreover, even when the control planarians of these two species did glide, the speed was significantly less than seen in *D. japonica* (Figure 4B-C) (p-values: 2.0 × 10^-28^, 6.2 × 10^-23^ [*S. mediterranea* Day 7 and 12] and 3.5 x10^-22^, 3.5 × 10^-12^ [*G. tigrina* Day 7 and 12]). In *S. mediterranea,* control gliding speeds were marginally greater than the resting speed cutoff of 0.3 mm/s. This value is extremely reduced compared to published *S. mediterranea* mean speeds of 1.62 mm/s (Talbot and Schötz, 2011) emphasizing the extreme lack of movement seen in these worms under these conditions.

**Figure 4.**
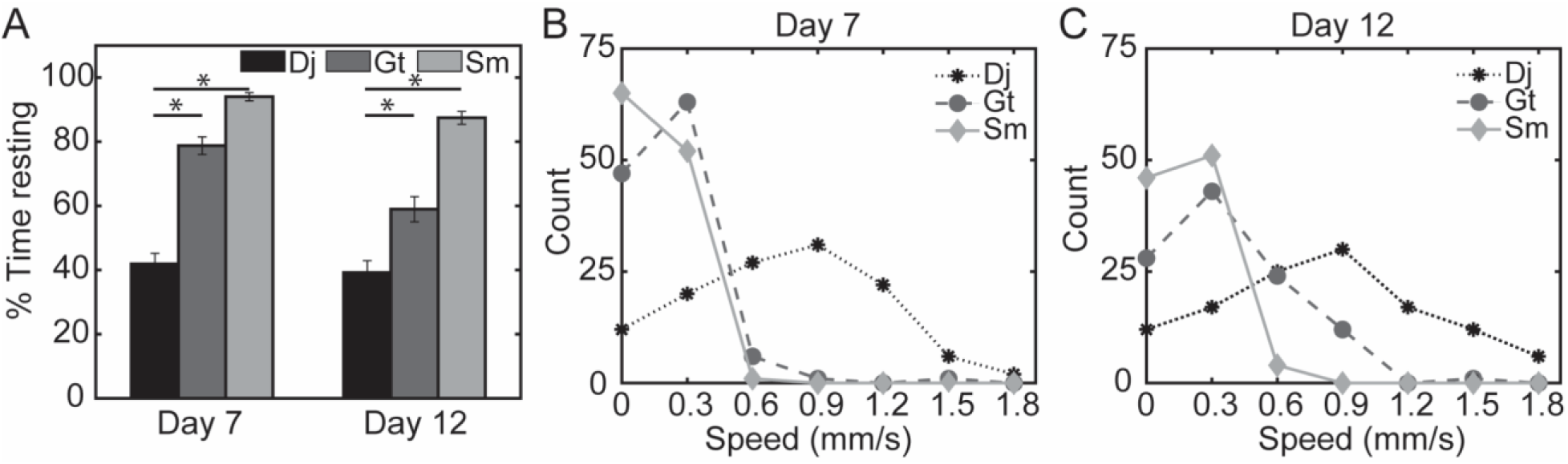
*S. mediterranea* and *G. tigrina* barely move during the locomotion assay. A) Average percent time spent resting in controls of different species on Days 7 and 12 during the unstimulated behavior assay. Error bars indicate ± SE. * indicates statistical significance with p-values < 0.05 as determined by a student’s t-test. B-C) Distribution of speeds of controls from different species at B) Day 7 or C) Day 12. For visualization, planarians which were resting for the entire assay were set to speeds of 0 mm/s.

To understand the reason for this difference in motility, we performed tests on intact *S. mediterranea.* Despite moving normally in a petri dish (Supplementary Figure 3A), these intact planarians exhibited reduced motility in the screening platform even when placed in 48-well plates within a few hours of petri dish testing. This motility defect was rescued if the planarians were imaged under bright white light (Supplementary Figure 3B), suggesting that the reason for *S. mediterranea* not to move in our assays on the screening platform was because the red lighting used for imaging is a wavelength that *S. mediterranea* are insensitive to (Paskin et al., 2014).

We also performed additional tests on *G. tigrina* planarians to investigate why their movement was reduced compared to *D. japonica.* Based on our previous data on population growth (Carter et al., 2015), we hypothesized that the small volume confinement of the 48-well plate may cause general health issues and increased immobility. To test this, we analyzed the performance of *G. tigrina* regenerating tails which were not stored in the 48-well plates but instead allowed to regenerate in petri dishes until screening. These planarians had no motility defects, whereas intact *G. tigrina* stored in sealed 48-well plates for 12 days displayed increased lethality and resting (Supplementary Figure 4). Together, these data suggest that *G. tigrina* move normally under the imaging conditions of the platform and that this population did not have general health issues but that the long-term storage conditions necessary for HTS (small volumes, sealed plate) are detrimental to *G. tigrina* health, leading to increased immobility in our screen. *S. mediterranea* motility was not significantly changed regardless of developmental condition (regenerating vs intact) or storage conditions (Supplementary Figure 5).

### Lack of motility hinders analysis of stimulated behaviors in S. mediterranea and G. tigrina

A major advantage of the planarian system is their complex repertoire of stereotypical behaviors in response to various stimuli, including light, temperature gradients, and noxious heat (Cochet-Escartin et al., 2015; Inoue et al., 2014, 2004; Paskin et al., 2014). We have found that our methodology for automated assessment of these behaviors in *D. japonica* is robust and sensitive to detect neuronal defects induced by neurotoxicants (Hagstrom et al., 2019; Zhang et al., 2019a, 2019b). Several of these behaviors (phototaxis and scrunching) have previously been evaluated in *S. mediterranea* and *G. tigrina* (Supplementary Table 1), albeit using low-throughput assays. Therefore, we evaluated whether these species were also capable of exhibiting robust stimulated behaviors using our automated methodology. Even though we found that *S. mediterranea* and *G. tigrina* showed decreased motility during unstimulated locomotion, it was still possible that the various stimuli could induce movement.

Planarians are negatively phototactic. Multiple planarian species have been shown to be most sensitive to blue light while being insensitive to red light (Davidson et al., 2011; Marriott, 1958; Paskin et al., 2014; Zhang et al., 2019a). Therefore, we exposed the planarians to 2 minutes of red light (dark cycle) followed by 1 minute of blue light (light cycle) and quantified the reaction as the difference in speeds between the light and dark cycles. Under these conditions, control *D. japonica* exhibited a robust increase in movement and speed during the light cycle, with an average speed difference of approximately 0.2 mm/s (Figure 5A-C). In contrast, *S. mediterranea* and *G. tigrina* control planarians exhibited much weaker reactions to the light, with average speed differences of only approximately 0.01-0.03 mm/s. These attenuated reactions were mainly a result of the immobility seen in these species as many of the planarians barely moved throughout the assay, regardless of the presence of the light (Figure 5A). As a result, the sensitivity of this assay in these species was limited.

**Figure 5.**
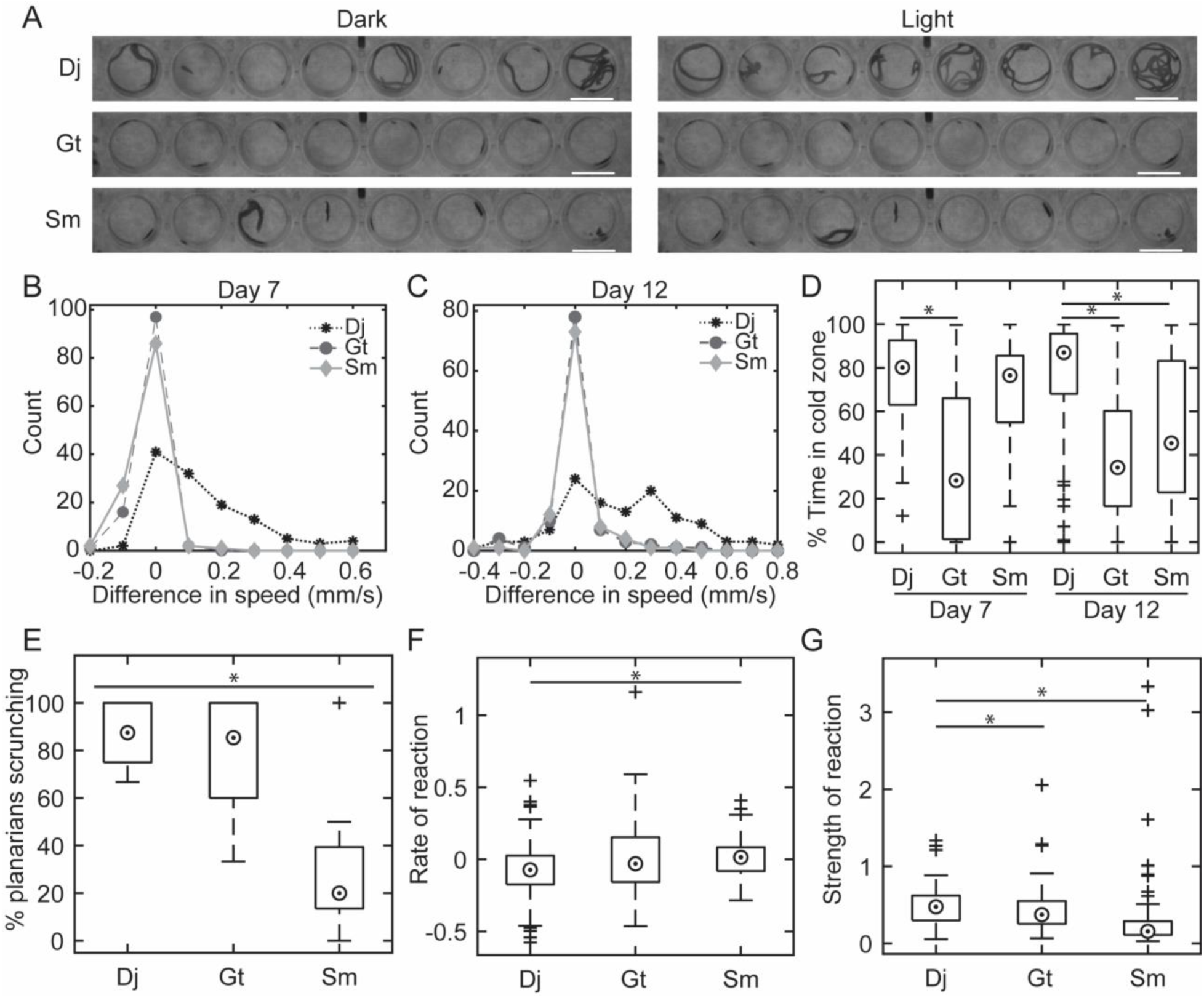
*S. mediterranea* and *G. tigrina* have attenuated performance in the various stimulated behavior assays. A) Minimum intensity projections showing the tracks of 8 control planarians in each species during the last minute of the dark cycle (Dark) or during the 1 minute blue light cycle (Light) in the phototaxis assay. Notice *D. japonica* planarians move more during the light period while the other two species barely move in either lighting. Scale bars: 10 mm. B-C) Distribution of the difference of average speed in the light and dark cycles during the phototaxis assay for control planarians of the different species at B) Day 7 or C) Day 12. The lower bin edge is plotted. D) Boxplot of the time spent in the cold zone during thermotaxis for controls of each species at Days 7 and 12. E) Boxplot of the percentage of control planarians scrunching in each replicate plate (n=8 per data point, n=15 data points per condition). F) Boxplot of the rate of reaction to noxious heat of controls for each species. G) Boxplot of the strength of reaction to noxious heat of controls for each species. For all boxplots, medians are shown as a dot in a circle, outliers are shown as crosses. For D-G, * indicates statistical significance with p-values < 0.05 as determined by a Mann-Whitney U-test.

Next, we evaluated how the different species performed in respect to thermotaxis. In this assay, a uniform temperature gradient is established within each well using a custom peltier setup, such that a cold region is established in each well, taking up an approximately 120 degree sector of the well (Zhang et al., 2019a). Control *D. japonica* prefer colder temperatures and thus spend the majority of time in the cold region (Figure 5D). In contrast, *G. tigrina* and *S. mediterranea* had significantly less robust preferences for the cold zone, especially at Day 12 (p-values: 1.1 × 10^-12^ [*G. tigrina*], 3.2 × 10^-6^ [*S. mediterranea*] compared to *D. japonica* using a Mann-Whitney U-Test) (Figure 5D). First, because this assay only uses data from moving worms, the sample size was greatly diminished in these two species due to the general immobility mentioned previously (Supplementary Table 2). Second, when the *S. mediterranea* and *G. tigrina* planarians did move, they spent less time in the cold region than *D. japonica* and had greater intraspecies variability (Figure 5D). Interestingly, *G. tigrina* reactions were similar to what would be expected from random motion across the well given that the cold sector is approximately 30% of the well area, suggesting these planarians do not react to the temperature gradient. *G. tigrina* planarians did not exhibit thermotaxis even when allowed to regenerate in petri dishes, and thus moved normally (Supplementary Figure 6A). Because we could neither induce thermotaxis in *G. tigrina* in the automated assay nor found any literature demonstrating thermotaxis in this species, we tested whether intact *G. tigrina* planarians could sense temperature gradients under low-throughput conditions using 6-well plates. In agreement with the HTS data, intact *G. tigrina* did not display thermotaxis under these conditions, while simultaneously assayed *D. japonica* planarians did. (Supplementary Figure 6B). These data suggest that *G. tigrina* do not exhibit thermotaxis in the same temperature ranges as the other two species.

Some of the *S. mediterranea* control planarians appeared to be successfully exhibiting thermotaxis and spent a majority of time in the cold region. However, the variability in this species was substantial and appears to be caused by their lack of motility. For example, *S. mediterranea* planarians that were resting near the cold zone showed successful thermotaxis, since they were able to sense the temperature gradient and move enough to enter the cold zone. To increase motility, we also tested thermotaxis under bright white light. Imaging under these conditions was sufficient to stimulate *S. mediterranea* to move (Supplementary Figure 3B) but failed to induce successful thermotaxis in *S. mediterranea* and *D. japonica* controls (Supplementary Figure 7). This suggests the addition of a light stimulus (bright white light) masked the behavioral response to the temperature gradient, in agreement with previous reports that when presented simultaneously, light is a stronger stimulus than temperature for *D. japonica* (Inoue et al., 2015).

Lastly, we evaluated the planarians’ ability to react to noxious heat. We have previously demonstrated that scrunching, a musculature-driven planarian escape gait that is conserved across species (Cochet-Escartin et al., 2015), can be induced by noxious heat (Cochet-Escartin et al., 2015; Sabry et al., 2019) and is a sensitive readout of neuronal function (Zhang et al., 2019a, 2019b). Scrunching was induced in approximately 88% of *D. japonica* control planarians under our experimental conditions; however, scrunching was much less prominent in the other two species (Figure 5E), with *S. mediterranea* showing a significantly lower scrunching induction rate compared to *D. japonica* (p-value< 1.8 × 10^-4^ using a Mann-Whitney U-Test).

In addition to binary classification of whether a planarian scrunched, we captured the dynamics of the noxious heat response by quantifying: 1) the rate at which the planarians responded to the heat and 2) the strength of their final reaction. (Supplementary Figure 1 and Materials and Methods). *S. mediterranea* and *G. tigrina* controls had weaker rates of reaction to the noxious heat compared to *D. japonica,* with a significant difference for *S. mediterranea* compared to *D. japonica* (p-value< 0.008; Mann-Whitney U-test) (Figure 5F). In *S. mediterranea* and *G. tigrina*, the median rate of reaction was approximately 0 (Figure 5F). This indicates little change in displacement, which results from the general lack of motility observed in these species (i.e. since these planarians were already not moving, a decrease in motion could not be assessed). Moreover, the lack of motility in these species greatly decreased the sample size for this endpoint (Supplementary Table 2). *G. tigrina* and *S. mediterranea* control planarians also showed significantly decreased “strength of reaction” scores compared to *D. japonica* (p-values: 0.03 [*G. tigrina*] and 4.7 × 10^-14^ [*S. mediterranea*]; Mann-Whitney U-test) (Figure 5G). These lower scores indicate these planarians were moving less than *D. japonica* during the second phase of the noxious heat assay, though this may be a result of their general lack of movement.

In summary, our data show that using our high-throughput methodology, motility and health issues in *S. mediterranea* and *G. tigrina* planarians greatly hindered our ability to assess the effects of chemical substances on morphology or behavior as even control animals demonstrated poor performance and high levels of variability.

### Toxicity of common solvents

The second aim of this study was to evaluate the effect of 4 common solvents in pharmacology and toxicology (DMSO, ethanol, methanol, ethyl acetate) on the three planarian species. Sorbitol served as a negative control. However, the lack of motility of *S. mediterranea* and *G. tigrina* planarians impaired our ability to evaluate solvent toxicity for certain endpoints due to data scarcity, as explained above. This effect was the greatest with *S. mediterranea*, resulting in many endpoints which could not be adequately evaluated (marked as “indeterminate” in Supplementary Figure 8). The only endpoint we could accurately use to compare solvent toxicity across all three species was lethality (Table 2). Of note, since we were interested in studying behavioral phenotypes in the absence of overt toxicity, the test concentrations had been chosen to not cause significant lethality in *D. japonica*.

**Table 2.** Lowest observed effect level for Day 12 lethality in each species. If lethality was not observed, the concentration is listed as > X, where X is the maximum tested concentration.

| Solvent | *D. japonica* | *G. tigrina* | *S. mediterranea* |
| --- | --- | --- | --- |
| DMSO | >1% | 1% | >1% |
| Ethanol | >1% | 0.05% | >1% |
| Methanol | >3.2% | 1.6% | 3.2% |
| Ethyl acetate | >0.04% | >0.04% | >0.04% |
| Sorbitol | >100 μM | >100 μM | >100 μM |

Overall, *D. japonica* planarians were less sensitive to the lethal effects of these solvents than the other two species. For methanol, lethality was observed in *G. tigrina* and *S. mediterranea* but not *D. japonica* at the concentrations tested (maximum 3.2%). *G. tigrina* showed the greatest sensitivity as lethality was observed in three of the tested solvents (DMSO, ethanol and methanol). The observed species differences in sensitivity highlight that care needs to be taken when extrapolating findings of chemical exposure between planarian species.

## Discussion and Conclusions

While existing studies have provided useful insight into how certain chemicals affect different aspects of planarian biology, the range of species and techniques used has made it difficult to compare results across different planarian studies to harmonize findings and contextualize how results in planarians relate to other species, especially humans.

Planarians and other new approach methods should have sufficiently high throughput to provide a robust, efficient alternative to existing testing methodologies. To this end, the use of multi-well plates and fully automated screening methodology is indispensable. Multi-well plates allow for the use of small testing volumes and the ability to test multiple conditions simultaneously, thus reducing chemical usage and experimental variability, respectively. We have found that, for *D. japonica*, 48-well plates provide a balance between throughput, maintaining planarian health long-term, and being able to robustly induce and quantify various behaviors. Moreover, in our testing paradigm, exposures are static with the plate sealed throughout exposure to reduce agitation to the planarians, reduce the amount of chemical required, and prevent changes in chemical concentration due to volatility/evaporation. Fully automated screening methodology is critical to obtain robust, unbiased results with sufficiently high throughput. Thus, we have focused our efforts on creating automated methodologies, in both the engineering of the screening platform and in the associated image and data analysis, such as for stickiness presented here. To ensure the necessary accuracy and robustness, all new automated analyses are manually cross-checked before full implementation, allowing us to refine the analysis as necessary.

Thus far, *D. japonica* is the only planarian species that has been successfully employed in large-scale automated screening (Zhang et al., 2019a, 2019b). Here, we have directly compared the performance of the 3 most commonly used freshwater planarian species in toxicology (*D. japonica*, *S. mediterranea,* and *G. tigrina)* under HTS conditions and evaluated their sensitivity to 4 common solvents (DMSO, ethanol, methanol and ethyl acetate). We found that *S. mediterranea* and *G. tigrina* are ill-suited for HTS because they do not display robust behaviors under the necessary experimental conditions (Figures 4-5, Supplementary Figures 3-4).

The reasons why the two species are not amenable to automated screening in 48-well plates differ between the two species. *S. mediterranea* exhibited limited locomotion when imaged with red light, but could be rescued using bright white light illumination (Supplementary Figure 3). This lack of motility prevented us from robustly evaluating locomotion or stimulated behaviors in the automated testing platform. Red lighting conditions are necessary to properly evaluate non-phototaxis behaviors, because the planarians’ response to light overrides other stimuli (Inoue et al., 2015) (Supplementary Figure 7). This lack of motility caused a major data loss for unstimulated locomotion and thermotaxis. In addition, *S. mediterranea* did not display a robust phototaxis response; it is unclear why, given that both *S. mediterranea* and *D. japonica* exhibit similar behaviors when exposed to a light gradient, though these behaviors often rely on moving planarians (Inoue et al., 2004; Paskin et al., 2014). Behavioral responses may also differ between exposure to a light gradient versus to a global light stimulus, as used here. While scrunching is one of the most sensitive readouts for assaying neurotoxicological effects in *D. japonica* (Zhang et al., 2019b, 2019a), we have been unable to robustly induce scrunching in *S. mediterranea* using a noxious heat bath here and in our previous work (Sabry et al., 2019). Together, these data suggest that *S. mediterranea* planarians are not well suited to multi-endpoint behavioral HTS. However, this species would be suitable to HTS assaying lethality, morphology and unstimulated behavior, if imaged using white light.

In contrast, *G. tigrina* moved normally under red light conditions when tested immediately after plate setup or if allowed to regenerate in petri dishes, but were negatively impacted by the confinement and small water volumes in the 48-well plates. Thus, while general health issues were not found in this species under normal laboratory conditions, their health declined over the 12 days of confinement, causing them to stop moving and/or die (Supplementary Figure 4), greatly limiting the number of planarians that could be analyzed (Supplementary Table 1). The observed health issues in *G. tigrina* in the small test volumes are perhaps not surprising since we have previously shown that *G. tigrina* are more sensitive to environmental conditions than *D. japonica* (Carter et al., 2015). Since *G. tigrina* planarians exhibit health problems during long-term storage in 48-well plates, this species is not suited for HTS of sub-chronic/chronic effects relying on small volume testing, independent of the details of the testing paradigm. Moreover, the lack of a thermotaxis response in *G. tigrina* without health or motility issues (Supplementary Figure 6) suggests this species may not have the same breadth of behaviors as *D. japonica*.

We have recently shown that both sensitivity and behavioral phenotypes to the pharmacological and toxicological effects of certain drugs can differ among *D. japonica* and *S. mediterranea* planarians (Sabry et al., 2019). Similarly, we have shown here that the 3 planarian species exhibit differential sensitivity to 4 common solvents. *G. tigrina* showed the greatest sensitivity to the tested solvents, though it is possible this sensitivity was a result of the general decline in health observed in this species under long-term confinement in 48-well plates. These species differences highlight that not all planarian research should be unified under a singular planarian model and that care needs to be taken when extrapolating from one planarian species to another. Moreover, the lower sensitivity of *D. japonica* planarians to these solvents suggests that higher solvent concentrations can be used in this species compared to the other two without fear of toxicological effects, further supporting our conclusion that this species is the best suited for toxicological research.

To be used in a regulatory context, new approach methods such as HTS in freshwater planarians must meet several “readiness criteria”, which evaluate the models’ technical capabilities, robustness, and relevancy to human health (Bal-Price et al., 2018; Crofton et al., 2011). A large aspect of this validation effort is to ensure results are reproducible across different laboratories. This necessitates that methods are transparent and standardized across different research groups. Our data here emphasize this need for method harmonization among planarian toxicological research as different species and different testing conditions produced significantly different effects. Our data show that, of the 3 most common planarian species used, only *D. japonica* is suitable for practical HTS conditions. We have also previously shown that data obtained with this species and our testing methodology is robust and relevant to mammalian outcomes (Hagstrom et al., 2019; Zhang et al., 2019b, 2019a). By standardizing testing methods, including the species used, the planarian toxicological community can work together towards validation of this promising invertebrate model.

## Acknowledgements

The authors thank Ziad Sabry for help with planarian care, and Dr. Bill Kristan for discussions and comments on the manuscript. Research reported in this publication was supported by the National Institute of Environmental Health Sciences of the National Institutes of Health under Award Number R15ES031354 (to E.M.S.C). The content is solely the responsibility of the authors and does not necessarily represent the official views of the National Institutes of Health.” V.B., D.C. and S. O. were funded through Swarthmore College summer fellowships.

## References

An, Y., Kawaguchi, A., Zhao, C., Toyoda, A., Sharifi-Zarchi, A., Mousavi, S.A., Bagherzadeh, R., Inoue, T., Ogino, H., Fujiyama, A., Chitsaz, H., Baharvand, H., Agata, K., 2018. Draft genome of Dugesia japonica provides insights into conserved regulatory elements of the brain restriction gene nou-darake in planarians. Zool. Lett. 4. https://doi.org/10.1186/s40851-018-0102-2

Bal-Price, A., Hogberg, H.T., Crofton, K.M., Daneshian, M., FitzGerald, R.E., Fritsche, E., Heinonen, T., Hougaard Bennekou, S., Klima, S., Piersma, A.H., Sachana, M., Shafer, T.J., Terron, A., Monnet-Tschudi, F., Viviani, B., Waldmann, T., Westerink, R.H.S., Wilks, M.F., Witters, H., Zurich, M.-G., Leist, M., 2018. Recommendation on test readiness criteria for new approach methods in toxicology: Exemplified for developmental neurotoxicity. ALTEX 35, 306–352. https://doi.org/10.14573/altex.1712081

Benazzi, M., 1993. Occurrence of a sexual population of Dugesia (Girardia) tigrina, a freshwater planarian native to America, in a lake of southern Italy. Ital. J. Zool. 60, 129–130. https://doi.org/10.1080/11250009309355799

Birkholz, T.R., Beane, W.S., 2017. The planarian TRPA1 homolog mediates extraocular behavioral responses to near-ultraviolet light. J. Exp. Biol. 220, 2616–2625. https://doi.org/10.1242/jeb.152298

Buttarelli, F.R., Pellicano, C., Pontieri, F.E., 2008. Neuropharmacology and behavior in planarians: Translations to mammals. Comp. Biochem. Physiol. - C Toxicol. Pharmacol. 147, 399–408. https://doi.org/10.1016/j.cbpc.2008.01.009

Byrne, T., 2018. Effects of ethanol on negative phototaxis and motility in brown planarians (Dugesia tigrina). Neurosci. Lett. 685, 102–108. https://doi.org/10.1016/j.neulet.2018.08.030

Carter, J.A., Lind, C.H., Truong, M.P., Collins, E.-M.S., 2015. To each his own. J. Stat. Phys. 161, 250–272. https://doi.org/10.1007/s10955-015-1310-1

Cebrià, F., Newmark, P.A., 2005. Planarian homologs of netrin and netrin receptor are required for proper regeneration of the central nervous system and the maintenance of nervous system architecture. Development 132, 3691–703. https://doi.org/10.1242/dev.01941

Cochet-Escartin, O., Mickolajczk, K.J., Collins, E.-M.S., 2015. Scrunching: a novel escape gait in planarians. Phys. Biol. 12, 055001. https://doi.org/doi: 10.1088/1478-3975/12/5/056010

Córdova López, A.M., Sarmento, R.A., de Souza Saraiva, A., Pereira, R.R., Soares, A.M.V.M., Pestana, J.L.T., 2019. Exposure to Roundup® affects behaviour, head regeneration and reproduction of the freshwater planarian Girardia tigrina. Sci. Total Environ. 675, 453–461. https://doi.org/10.1016/j.scitotenv.2019.04.234

Crofton, K.M., Mundy, W.R., Lein, P.J., Bal-Price, A., Coecke, S., Seiler, A.E.M., Knaut, H., Buzanska, L., Goldberg, A., 2011. Developmental neurotoxicity testing: recommendations for developing alternative methods for the screening and prioritization of chemicals. ALTEX 28, 9–15.

Currie, K.W., Pearson, B.J., 2013. Transcription factors lhx1/5-1 and pitx are required for the maintenance and regeneration of serotonergic neurons in planarians. Development 140, 3577–88. https://doi.org/10.1242/dev.098590

Davidson, C., Prados, J., Gibson, C.L., Young, A.M.J., Barnes, D., Sherlock, R., Hutchinson, C. V., 2011. Shedding light on photosensitive behaviour in brown planaria (Dugesia Tigrina). Perception 40, 743–746. https://doi.org/10.1068/p6949

Dunkel, J., Talbot, J., Schötz, E.-M., 2011. Memory and obesity affect the population dynamics of asexual freshwater planarians. Phys. Biol. 8, 026003. https://doi.org/10.1088/1478-3975/8/2/026003

Fritsche, E., Grandjean, P., Crofton, K.M., Aschner, M., Goldberg, A., Heinonen, T., Hessel, E.V.S.S., Hogberg, H.T., Bennekou, S.H., Lein, P.J., Leist, M., Mundy, W.R., Paparella, M., Piersma, A.H., Sachana, M., Schmuck, G., Solecki, R., Terron, A., Monnet-Tschudi, F., Wilks, M.F., Witters, H., Zurich, M.-G.G., Bal-Price, A., 2018. Consensus statement on the need for innovation, transition and implementation of developmental neurotoxicity (DNT) testing for regulatory purposes. Toxicol. Appl. Pharmacol. 354, 3–6.

Garcia-Fernandez, J., Bayascas-Ramirez, J.R., Marfany, G., Munoz-Marmol, A.M., Casali, A., Baguna, J., Salo, E., 1995. High copy number of highly similar mariner-like transposons in planarian (Platyhelminthe): evidence for a trans-phyla horizontal transfer. Mol. Biol. Evol. 12, 421–431. https://doi.org/10.1093/oxfordjournals.molbev.a040217

Grohme, M.A., Schloissnig, S., Rozanski, A., Pippel, M., Young, G.R., Winkler, S., Brandl, H., Henry, I., Dahl, A., Powell, S., Hiller, M., Myers, E., Rink, J.C., 2018. The genome of Schmidtea mediterranea and the evolution of core cellular mechanisms. Nature 554, 56–61. https://doi.org/10.1038/nature25473

Hagstrom, D., Cochet-Escartin, O., Collins, E.-M.S., 2016. Planarian brain regeneration as a model system for developmental neurotoxicology. Regeneration 3, 65–77. https://doi.org/10.1002/reg2.52

Hagstrom, D., Cochet-Escartin, O., Zhang, S., Khuu, C., Collins, E.-M.S., 2015. Freshwater planarians as an alternative animal model for neurotoxicology. Toxicol. Sci. 147, 270–285. https://doi.org/10.1093/toxsci/kfv129

Hagstrom, D., Truong, L., Zhang, S., Tanguay, R., Collins, E.-M.S., 2019. Comparative analysis of zebrafish and planarian model systems for developmental neurotoxicity screens using an 87-compound library. Toxicol. Sci. 167, 15–25. https://doi.org/10.1093/toxsci/kfy180

Hagstrom, D., Zhang, S., Ho, A., Tsai, E.S., Radić, Z., Jahromi, A., Kaj, K.J., He, Y., Taylor, P., Collins, E.M.S., 2018. Planarian cholinesterase: molecular and functional characterization of an evolutionarily ancient enzyme to study organophosphorus pesticide toxicity. Arch. Toxicol. 92, 1161–1176. https://doi.org/10.1007/s00204-017-2130-7

Hoshino, K., Ohnishi, K., Yoshida, W., Shinozawa, T., 1991. Analysis of ploidy in a planarian by flow cytometry. Hydrobiologia 227, 175–178. https://doi.org/10.1007/BF00027599

Inoue, T., Hoshino, H., Yamashita, T., Shimoyama, S., Agata, K., 2015. Planarian shows decision-making behavior in response to multiple stimuli by integrative brain function. Zool. Lett. 1. https://doi.org/10.1186/s40851-014-0010-z

Inoue, T., Kumamoto, H., Okamoto, K., Umesono, Y., Sakai, M., Alvarado, A.S., Agata, K., 2004. Morphological and functional recovery of the planarian photosensing system during head regeneration. Zoolog. Sci. 21, 275–283. https://doi.org/10.2108/zsj.21.275

Inoue, T., Yamashita, T., Agata, K., 2014. Thermosensory signaling by TRPM is processed by brain serotonergic neurons to produce planarian thermotaxis. J. Neurosci. 34, 15701–14. https://doi.org/10.1523/JNEUROSCI.5379-13.2014

Knakievicz, T., Ferreira, H.B., 2008. Evaluation of copper effects upon Girardia tigrina freshwater planarians based on a set of biomarkers. Chemosphere 71, 419–28.

Lein, P., Silbergeld, E., Locke, P., Goldberg, A.M., 2005. In vitro and other alternative approaches to developmental neurotoxicity testing (DNT). Environ. Toxicol. Pharmacol. 19, 735–44. https://doi.org/10.1016/j.etap.2004.12.035

Li, C.H., Tam, P.K.S., 1998. An iterative algorithm for minimum cross entropy thresholding. Pattern Recognit. Lett. 19, 771–776. https://doi.org/10.1016/S0167-8655(98)00057-9

Lowe, J.R., Mahool, T.D., Staehle, M.M., 2015. Ethanol exposure induces a delay in the reacquisition of function during head regeneration in Schmidtea mediterranea. Neurotoxicol. Teratol. 48, 28–32.

Malinowski, P.T., Cochet-Escartin, O., Kaj, K.J., Ronan, E., Groisman, A., Diamond, P.H., Collins, E.-M.S., 2017. Mechanics dictate where and how freshwater planarians fission. Proc. Natl. Acad. Sci. U. S. A. 114, 10888–10893. https://doi.org/10.1073/pnas.1700762114

Marriott, F.H.C., 1958. The absolute light-sensitivity and spectral threshold curve of the aquatic flatworm Dendrocoelum lacteum. J. Physiol. 143, 369–379. https://doi.org/10.1113/jphysiol.1958.sp006065

Martin, G.G., 1978. A new function of rhabdites: Mucus production for ciliary gliding. Zoomorphologie 91, 235–248. https://doi.org/10.1007/BF00999813

Mineta, K., Nakazawa, M., Cebria, F., Ikeo, K., Agata, K., Gojobori, T., 2003. Origin and evolutionary process of the CNS elucidated by comparative genomics analysis of planarian ESTs. Proc. Natl. Acad. Sci. U. S. A. 100, 7666–71.

Moustakas, D., Mezzio, M., Rodriguez, B.R., Constable, M.A., Mulligan, M.E., Voura, E.B., 2015. Guarana provides additional stimulation over caffeine alone in the planarian model. PLoS One 10, e0123310. https://doi.org/10.1371/journal.pone.0123310

Nishimura, K., Kitamura, Y., Taniguchi, T., Agata, K., 2010. Analysis of motor function modulated by cholinergic neurons in planarian Dugesia japonica. Neuroscience 168, 18–30. https://doi.org/10.1016/j.neuroscience.2010.03.038

Nishimura, K., Kitamura, Y., Umesono, Y., Takeuchi, K., Takata, K., Taniguchi, T., Agata, K., 2008. Identification of glutamic acid decarboxylase gene and distribution of GABAergic nervous system in the planarian Dugesia japonica. Neuroscience 153, 1103–14. https://doi.org/10.1016/j.neuroscience.2008.03.026

Otsu, N., 1979. Threshold selection method from gray-level histograms. IEEE Trans Syst Man Cybern SMC-9, 62–66. https://doi.org/10.1109/tsmc.1979.4310076

Paskin, T.R., Jellies, J., Bacher, J., Beane, W.S., 2014. Planarian phototactic assay reveals differential behavioral responses based on wavelength. PLoS One 9, e114708. https://doi.org/10.1371/journal.pone.0114708

Passarelli, F., Merante, A., Pontieri, F.E., Margotta, V., Venturini, G., Palladini, G., 1999. Opioid-dopamine interaction in planaria: a behavioral study. Comp. Biochem. Physiol. C. Pharmacol. Toxicol. Endocrinol. 124, 51–5.

Pearce, R.G., Setzer, R.W., Strope, C.L., Sipes, N.S., Wambaugh, J.F., 2017. httk : R package for high-throughput toxicokinetics. J. Stat. Softw. 79. https://doi.org/10.18637/jss.v079.i04

Plusquin, M., Stevens, A.-S., Van Belleghem, F., Degheselle, O., Van Roten, A., Vroonen, J., Blust, R., Cuypers, A., Artois, T., Smeets, K., 2012. Physiological and molecular characterisation of cadmium stress in Schmidtea mediterranea. Int. J. Dev. Biol. 56, 183–91.

Poirier, L., Brun, L., Jacquet, P., Lepolard, C., Armstrong, N., Torre, C., Daudé, D., Ghigo, E., Chabrière, E., 2017. Enzymatic degradation of organophosphorus insecticides decreases toxicity in planarians and enhances survival. Sci. Rep. 7, 15194. https://doi.org/10.1038/s41598-017-15209-8

Ramakrishnan, L., DeSaer, C., 2011. Carbamazepine inhibits distinct chemoconvulsant-induced seizure-like activity in Dugesia tigrina. Pharmacol. Biochem. Behav. 99, 665–670. https://doi.org/10.1016/j.pbb.2011.06.003

Robb, S.M.C., Ross, E., Sánchez Alvarado, A., 2008. SmedGD: the Schmidtea mediterranea genome database. Nucleic Acids Res. 36, D599–606.

Rompolas, P., Patel-King, R.S., King, S.M., 2010. An outer arm Dynein conformational switch is required for metachronal synchrony of motile cilia in planaria. Mol. Biol. Cell 21, 3669– 79. https://doi.org/10.1091/mbc.E10-04-0373

Ross, K.G., Currie, K.W., Pearson, B.J., Zayas, R.M., 2017. Nervous system development and regeneration in freshwater planarians. Wiley Interdiscip. Rev. Dev. Biol. 6, e266. https://doi.org/10.1002/wdev.266

Rozanski, A., Moon, H., Brandl, H., Martín-Durán, J.M., Grohme, M.A., Hüttner, K., Bartscherer, K., Henry, I., Rink, J.C., 2019. PlanMine 3.0—improvements to a mineable resource of flatworm biology and biodiversity. Nucleic Acids Res. 47, D812–D820. https://doi.org/10.1093/nar/gky1070

Sabry, Z., Ho, A., Ireland, D., Rabeler, C., Cochet-Escartin, O., Collins, E.M.S., 2019. Pharmacological or genetic targeting of Transient Receptor Potential (TRP) channels can disrupt the planarian escape response. PLoS One 14, e0226104. https://doi.org/10.1371/journal.pone.0226104

Stevens, A.S., Pirotte, N., Plusquin, M., Willems, M., Neyens, T., Artois, T., Smeets, K., 2014. Toxicity profiles and solvent-toxicant interference in the planarian Schmidtea mediterranea after dimethylsulfoxide (DMSO) exposure. J. Appl. Toxicol. 35, 319–326. https://doi.org/10.1002/jat.3011

Talbot, J., Schötz, E.-M., 2011. Quantitative characterization of planarian wild-type behavior as a platform for screening locomotion phenotypes. J. Exp. Biol. 214, 1063–7. https://doi.org/10.1242/jeb.052290

Thomas, R.S., Bahadori, T., Buckley, T.J., Cowden, J., Deisenroth, C., Dionisio, K.L., Frithsen, J.B., Grulke, C.M., Gwinn, M.R., Harrill, J.A., Higuchi, M., Houck, K.A., Hughes, M.F., Hunter, E.S., Isaacs, K.K., Judson, R.S., Knudsen, T.B., Lambert, J.C., Linnenbrink, M., Martin, T.M., Newton, S.R., Padilla, S., Patlewicz, G., Paul-Friedman, K., Phillips, K.A., Richard, A.M., Sams, R., Shafer, T.J., Setzer, R.W., Shah, I., Simmons, J.E., Simmons, S.O., Singh, A., Sobus, J.R., Strynar, M., Swank, A., Tornero-Valez, R., Ulrich, E.M., Villeneuve, D.L., Wambaugh, J.F., Wetmore, B.A., Williams, A.J., 2019. The next generation blueprint of computational toxicology at the U.S. Environmental Protection Agency. Toxicol. Sci. 169, 317–332. https://doi.org/10.1093/toxsci/kfz058

Tran, T.A., Hesler, M., Moriones, O.H., Jimeno-Romero, A., Fischer, B., Bastús, N.G., Puntes, V., Wagner, S., Kohl, Y.L., Gentile, L., 2019. Assessment of iron oxide nanoparticle ecotoxicity on regeneration and homeostasis in the replacement model system Schmidtea mediterranea. ALTEX 36, 583–596. https://doi.org/10.14573/altex.1902061

Tsuji, R., Crofton, K.M., 2012. Developmental neurotoxicity guideline study: Issues with methodology, evaluation and regulation. Congenit. Anom. (Kyoto). 52, 122–128. https://doi.org/10.1111/j.1741-4520.2012.00374.x

US EPA, 2018. Strategic Plan to Promote the Development and Implementation of Alternative Test Methods Within the TSCA Program. Washington, D.C.

Van Der Walt, S., Schönberger, J.L., Nunez-Iglesias, J., Boulogne, F., Warner, J.D., Yager, N., Gouillart, E., Yu, T., 2014. Scikit-image: Image processing in python. PeerJ 2014. https://doi.org/10.7717/peerj.453

Wheeler, A.R., 2019. Directive to Prioritize Efforts to Reduce Animal Testing. Washington, D.C.

Wheeler, N.J., Agbedanu, P.N., Kimber, M.J., Ribeiro, P., Day, T.A., Zamanian, M., 2015. Functional analysis of Girardia tigrina transcriptome seeds pipeline for anthelmintic target discovery. Parasit. Vectors 8, 34. https://doi.org/10.1186/s13071-014-0622-3

Wu, J.P., Li, M.H., 2018. The use of freshwater planarians in environmental toxicology studies: Advantages and potential. Ecotoxicol. Environ. Saf. https://doi.org/10.1016/j.ecoenv.2018.05.057

Zhang, S., Hagstrom, D., Hayes, P., Graham, A., Collins, E.-M.S., 2019a. Multi-behavioral endpoint testing of an 87-chemical compound library in freshwater planarians. Toxicol. Sci. 167, 26–44. https://doi.org/10.1093/toxsci/kfy145

Zhang, S., Ireland, D., Sipes, N.S., Behl, M., Collins, E.-M.S., 2019b. Screening for neurotoxic potential of 15 flame retardants using freshwater planarians. Neurotoxicol. Teratol. 73, 54– 66. https://doi.org/10.1016/j.ntt.2019.03.003

