## Supplementary Materials for "*Dugesia japonica* is the best suited of three planarian species for high-throughput toxicology screening"

### Supplementary Tables

**Supplementary Table 1. Overview of qualitative and quantitative behavioral assays in *D. japonica, S. mediterranea,* and *G. tigrina***

| Behavior | Species | Throughput[1] | Analysis | References |
|---|---|---|---|---|
| Locomotion | Dj | Low, high | pLMV[2], automated tracking | (Hagstrom et al., 2015; Lambrus et al., 2015; Zhang et al., 2019) |
| | Sm | Low | pLMV, automated tracking | (Lowe et al., 2015; Plusquin et al., 2012; Stevens et al., 2014; Talbot and Schötz, 2011) |
| | Gt | Low | pLMV | (Knakievicz and Ferreira, 2008; Moustakas et al., 2015) |
| Phototaxis | Dj | Low, high | Automated tracking | (Balestrini et al., 2014; Inoue et al., 2004; Zhang et al., 2019) |
| | Sm | Low | Quadrant location, automated tracking | (Lambrus et al., 2015; Lowe et al., 2015; Paskin et al., 2014) |
| | Gt | Low | Gridlines crossed | (Byrne, 2018) |
| Thermotaxis | Dj | Low, high | Automated tracking | (Hagstrom et al., 2015; Inoue et al., 2014; Zhang et al., 2019) |
| Scrunching | Dj | Low, high | Automated tracking | (Cochet-Escartin et al., 2015; Sabry et al., 2019; Zhang et al., 2019) |
| | Sm | Low, high | Automated tracking | (Cochet-Escartin et al., 2015; Sabry et al., 2019) |
| | Gt | Low | Automated tracking | (Cochet-Escartin et al., 2015) |
| Chemotaxis | Dj | Low | Quadrant location | (Inoue et al., 2015) |
| Rheosensation | Sm | Low | Manual image analysis | (Ross et al., 2018) |
| Vibriosensation | Sm | Low | Automated tracking | (Ross et al., 2018) |
| Noxious heat response | Dj | Low | Automated tracking | (Hagstrom et al., 2018) |
| | Sm | Low | Automated tracking | (Arenas et al., 2017) |
| Thigmotaxis | Dj | Low | Quadrant location | (Inoue et al., 2015) |

Dj: *Dugesia japonica*; Sm: *Schmidtea mediterranea*; Gt: *Girardia tigrina*

[1] Throughput was marked as “low” for assays which tested planarians individually or in bulk culture or “high” when multiple planarians were assayed simultaneously, such as in multi-well plates.
[2] pLMV (planarian locomotor velocity) as first described in (Raffa et al., 2001).

**Supplementary Table 2. Number of total analyzable controls for each endpoint and species.** Maximum possible n=120.

| **Endpoint** | ***D. japonica*** | ***G. tigrina*** | ***S. mediterranea*** |
|---|---|---|---|
| Lethality – Day 7 | 120 | 120 | 120 |
| Lethality – Day 12 | 120 | 120 | 109 |
| Body shape - Day 7 | 120 | 115 | 116 |
| Body shape - Day 12 | 119 | 106 | 95 |
| Eye regeneration – Day 7 | 104 | 43 | 57 |
| Stickiness – Day 7 | 117 | 104 | 111 |
| Stickiness – Day 12 | 118 | 88 | 76 |
| % time resting – Day 7 | 120 | 118 | 118 |
| % time resting – Day 12 | 119 | 108 | 101 |
| Speed – Day 7 | 108 | 71 | 53 |
| Speed – Day 12 | 107 | 80 | 55 |
| Phototaxis – Day 7 | 120 | 118 | 118 |
| Phototaxis – Day 12 | 119 | 108 | 101 |
| Thermotaxis - Day 7 | 101 | 22 | 43 |
| Thermotaxis - Day 12 | 108 | 52 | 38 |
| Scrunching | 117 | 88 | 74 |
| Noxious stimuli - rate | 111 | 45 | 28 |
| Noxious stimuli - strength | 118 | 102 | 95 |

Supplementary Figures

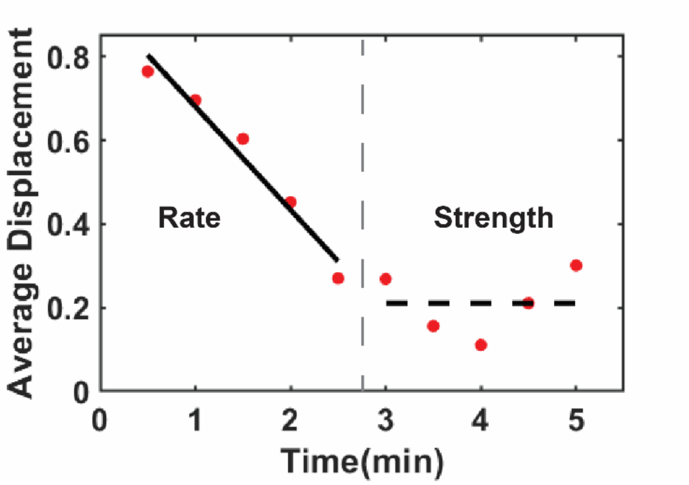


**Supplementary Figure 1. Quantification of the rate and strength of reaction to noxious heat.** To quantify the dynamics of a planarian's response to noxious heat, the displacement across 6 second intervals were calculated and then averaged across 30 second bins (red dots). The rate of reaction was quantified as the slope of the average displacements in the first 2.5 minutes of the assay (solid line). The strength of the reaction was quantified as the mean of the average displacements during minutes 3-5 (black dashed line). Data from one representative *D. japonica* planarian are shown.

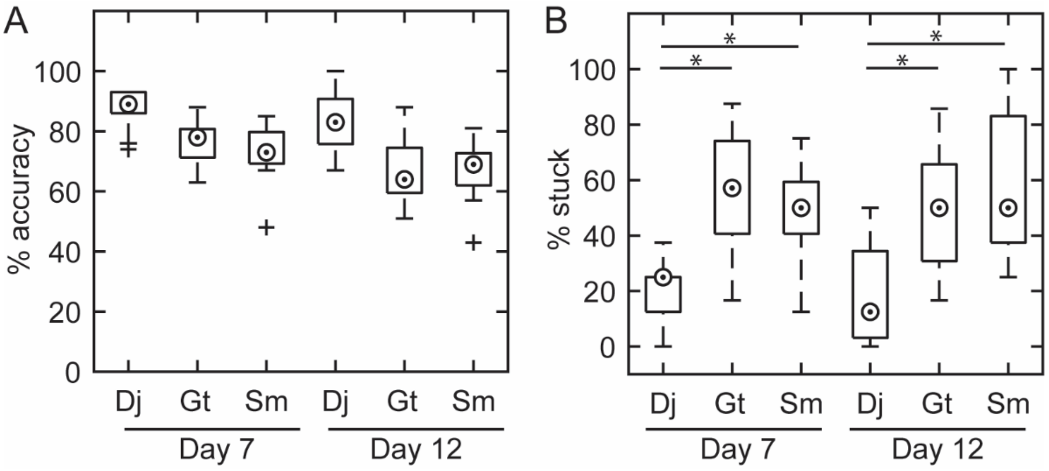


**Supplementary Figure 2. Evaluation of automated stickiness analysis.** A) Boxplot of the accuracy of the automated stickiness analysis for each species in each replicate plate. (n=8 per data point, n=15 data points per condition). B) Boxplot of the percent control worms stuck in each replicate plate (n=8 per data point, n=15 data points per condition) as determined by the automated analysis. Medians are shown as a dot in a circle, outliers are shown as crosses for A-B. * indicates statistical significance with p-values < 0.05 using the Mann-Whitney U-Test. The same trends in relative stickiness across species were found with both manual and automated analysis (compare to Figure 3C).

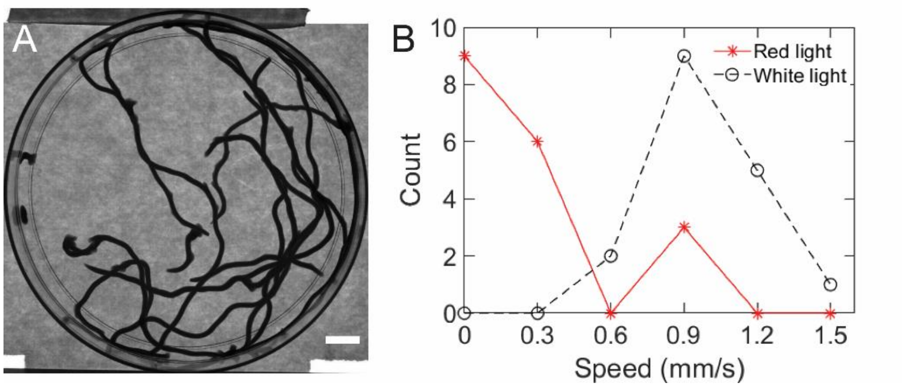


**Supplementary Figure 3. *S. mediterranea* motility defect is due to imaging in red light.** A) Minimum intensity projection showing intact *S. mediterranea* (n=16) were able to move normally in a petri dish before loading into the 48-well plate. B) Distribution of speeds in intact *S. mediterranea* planarians imaged under unstimulated behavioral assay conditions in 48-well plates under red or white light. For visualization, planarians which were resting for the entire assay had their speeds set to 0 mm/s. The lower limit of each bin edge is plotted.

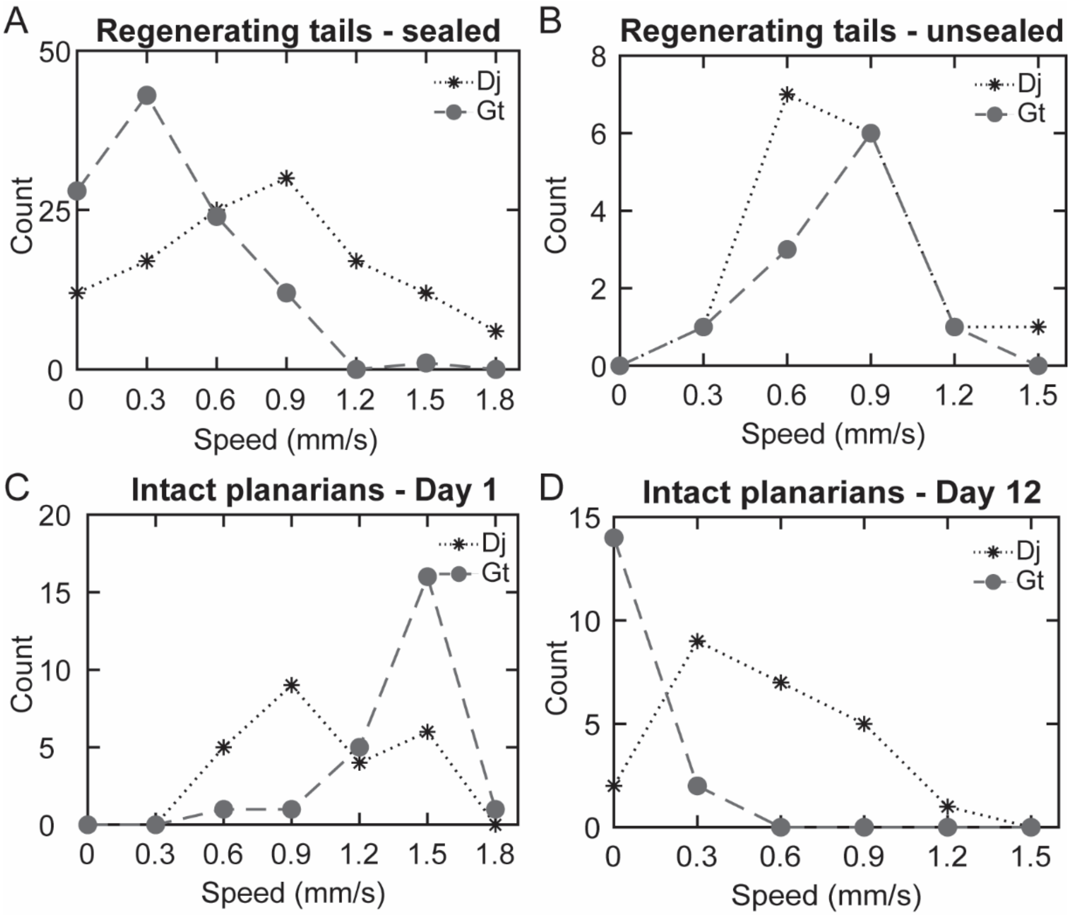


**Supplementary Figure 4.** ***G. tigrina*** **motility defect is due to long-term storage in sealed 48-well plates.** A-B) Distribution of speeds in control/wild-type Day 12 regenerating *D. japonica* (Dj) and *G. tigrina* (Gt) planarians which were kept either in A) sealed 48-well plates or B) petri dishes between screening days. Data from A are the same as presented in Figure 4C. C-D) Distribution of speeds in wild-type *D. japonica* and *G. tigrina* intact planarians on C) Day 1 or D) after being kept in sealed 48-well plates for 12 days. For D, 8 of the 24 *G. tigrina* planarians initially placed in the screening plate died by Day 12. No deaths were seen in *D. japonica*. For visualization, planarians which were resting for the entire assay had their speeds set to 0 mm/s. The lower limit of each bin edge is plotted.

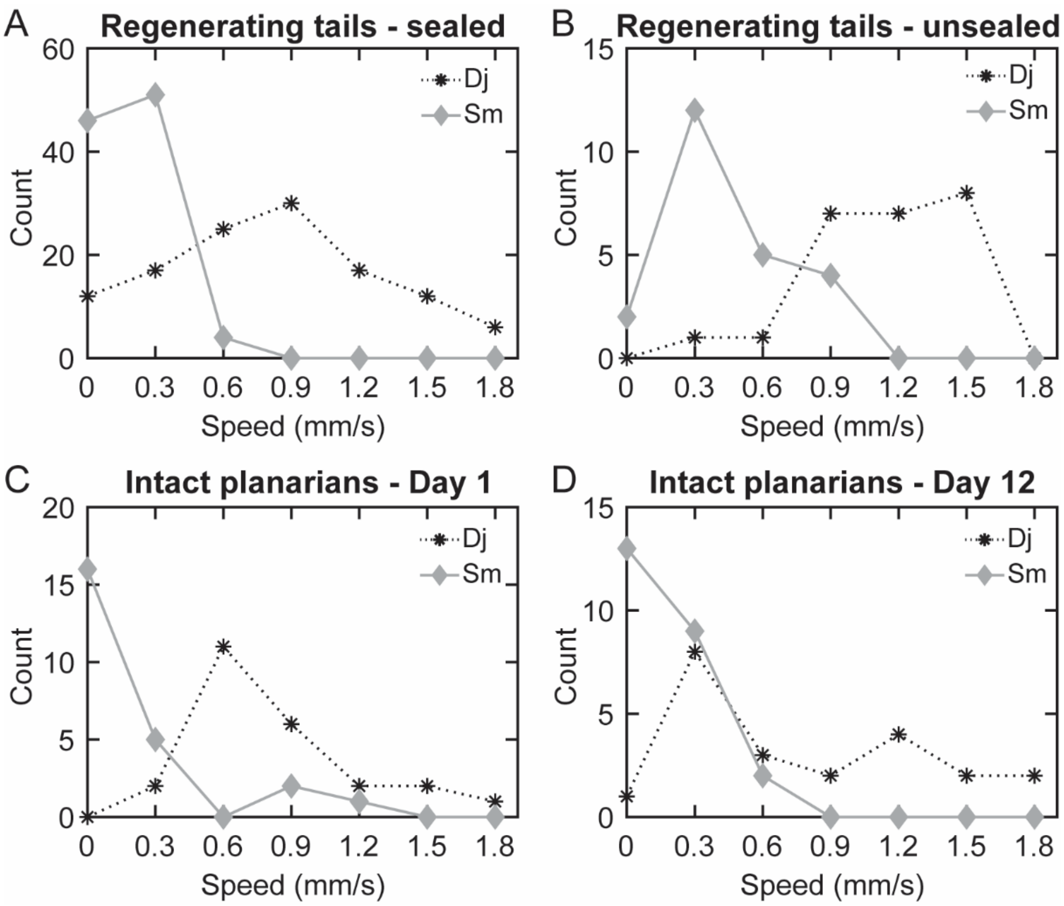


**Supplementary Figure 5.** ***S. mediterranea*** **motility is not affected by long-term storage in sealed 48-well plates.** A-B) Distribution of speeds in control/wild-type Day 12 regenerating *D. japonica* (Dj) and *S. mediterranea* (Sm) planarians which were kept either in A) sealed 48-well plates or B) petri dishes between screening days. Data from A are the same as presented in Figure 4C. C-D) Distribution of speeds in wild-type *D. japonica* and *S. mediterranea* intact planarians on C) Day 1 or D) after being kept in sealed 48-well plates for 12 days. For visualization, planarians which were resting for the entire assay had their speeds set to 0 mm/s. The lower limit of each bin edge is plotted.

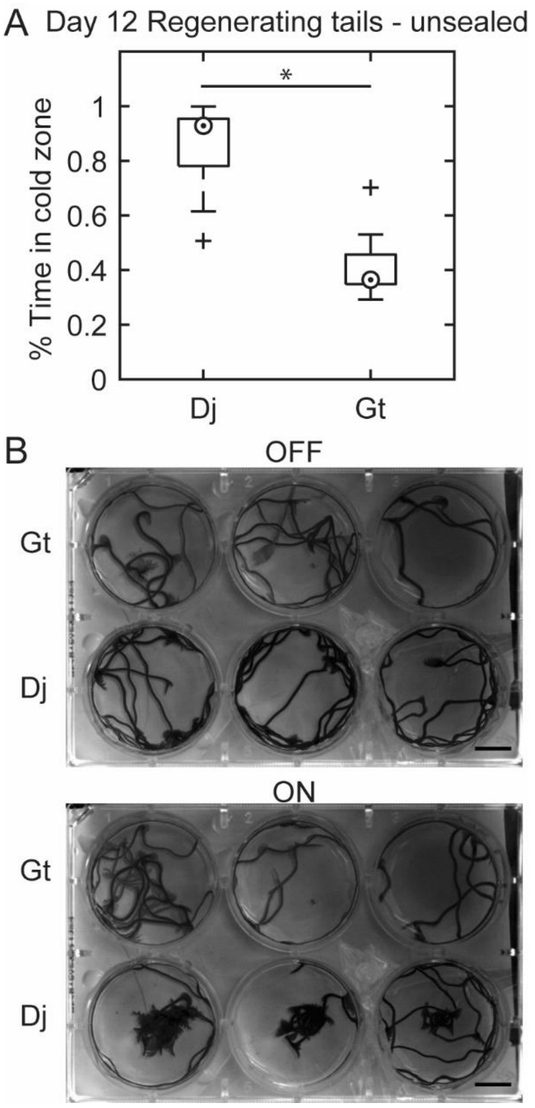


**Supplementary Figure 6. *G. tigrina* do not perform thermotaxis.** A) Boxplot of the time spent in the cold zone during thermotaxis for control *D. japonica* (Dj, n=15) and *G. tigrina* (Gt, n=10) regenerating planarians allowed to regenerate in petri dishes. Note these are the same planarians as tested in Supplementary Figure 4B. Medians are shown as a dot in a circle, outliers are shown as crosses. * indicates statistical significance with p-values $< 0.05$ as determined by a Mann-Whitney U-test. B) Minimum intensity projections of wild-type intact *G. tigrina* and *D. japonica* planarians with the peltier off (top) or on (bottom). Note that when the peltier is on, the *D. japonica* planarians spend the most time in the cold region in the middle of the well, while *G. tigrina* planarians continue to move randomly across the well. Scale bars: 10 mm. Recording duration: 2 min for each (OFF/ON). Images are representative of n=3 independent experiments.

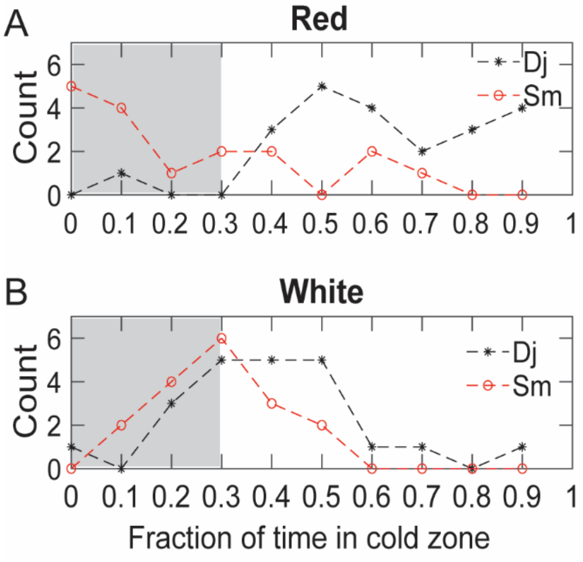


**Supplementary Figure 7. *D. japonica* and *S. mediterranea* do not exhibit thermotaxis in white light.** A-B) Distribution of the fraction of time spent in the cold zone for intact wild-type *D. japonica* and *S. mediterranea* when imaged under A) red light (our normal conditions) or B) white light. The lower limit of each bin edge is plotted. Since the cold zone is approximately 1/3 of the well, fractions of 0.3 and lower (in gray) likely represent random motion across the well.

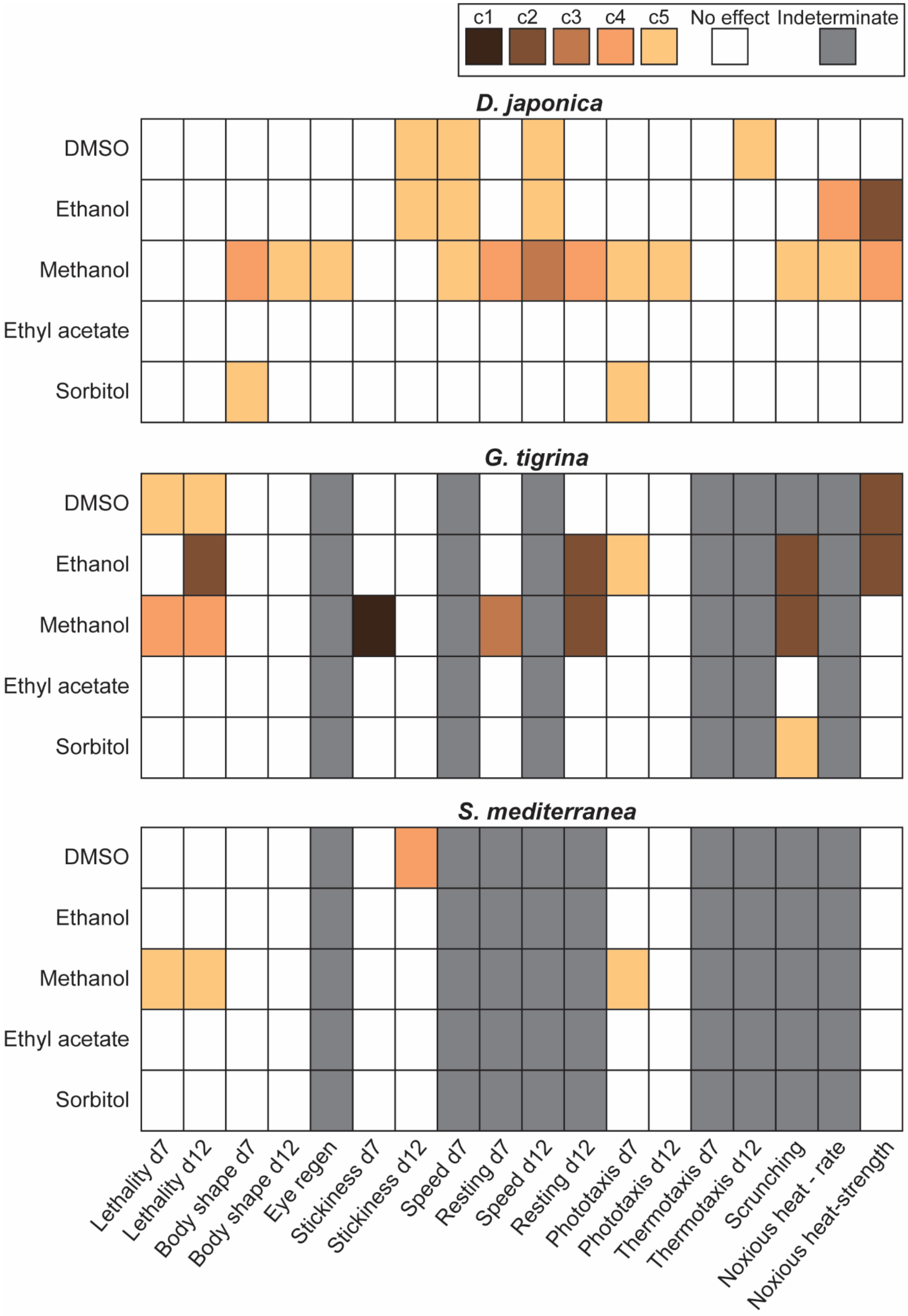
c1
c2
c3
c4
c5
No effect
Indeterminate
D. japonica
G. tigrina
S. mediterranea
DMSO
Ethanol
Methanol
Ethyl acetate
Sorbitol
Lethality d7
Lethality d12
Body shape d7
Body shape d12
Eye regen
Stickiness d7
Stickiness d12
Speed d7
Resting d7
Speed d12
Resting d12
Phototaxis d7
Phototaxis d12
Thermotaxis d7
Thermotaxis d12
Scrunching
Noxious heat - rate
Noxious heat-strength

**Supplementary Figure 8. Summary of screening data.** Heatmaps of the lowest concentration which elicited a statistically significant effect for each endpoint, chemical, and species. Only concentration-dependent hits are shown. Tested concentrations are shown as c1 (lowest) to c5 (highest), see Table 1 for concentrations. Chemicals which did not elicit a statistically significant effect in any of the tested concentrations are shown as "no effect". Endpoints which could not be evaluated due to low sample sizes or large control variability are shown as "indeterminate". d7: Day 7, d12: Day 12.
